# The natural flavonoid dihydromyricetin targets senescent cells *via* PRDX2 and alleviates age-related diseases

**DOI:** 10.64898/2025.12.25.696450

**Authors:** Qixia Xu, Gaoxiang Li, Hongwei Zhang, Zhirui Jiang, Xiuxia Gao, Zi Li, Larissa G. P. Langhi Prata, James L. Kirkland, Guilong Zhang, Yu Sun

## Abstract

Aging is a preeminent risk factor for chronic diseases, with cellular senescence as one of the major hallmarks and an effective target to delay, prevent or alleviate age-related disorders. Here we report *in vitro* screening outputs from a natural medicinal agent (NMA) library, wherein dihydromyricetin (DMY), a natural flavonoid, showed senotherapeutic potential. DMY protected senescent fibroblasts from further DNA damage and attenuated the senescence-associated secretory phenotype (SASP), acting as a senomorphic agent. Proteomics suggested that DMY promotes nuclear translocation of peroxiredoxin 2 (PRDX2), a reactive oxygen species (ROS) scavenger, facilitating DNA repair in senescent cells. In prematurely aged mice, DMY administration mitigated tissue aging changes and age-related physiological decline. In anticancer regimens, DMY improved outcomes of chemotherapy. However, DMY demonstrated senolytic activity against senescent microglial cells, wherein basal PRDX2 expression remains low, by impairing mitochondrial function to promote ROS-mediated apoptosis. In mice developing an Alzheimer’s disease (AD)-like state, DMY eliminated senescent microglial cells from amyloid β-protein (Aβ) plaques, alleviating Aβ-associated pathological phenotypes. Together, our study proposes DMY to be a natural senotherapeutic agent representing a therapeutic solution for mitigating age-related morbidities including but not limited to cancers and AD.

## Introduction

Aging is characterized by time-dependent declines in the functional integrity of multiple organs, which underly the pathogenesis of multiple age-related disorders ^1^. Insights into mechanisms that allow aging to promote diseases are critical for effectively combating adverse consequences of aging. As a hallmark of organismal aging, cellular senescence appears to have evolved to have protective functions *in vivo*, occurring in response to macromolecular damage, including those caused by replicative exhaustion (replicative senescence, RS), oncogenic activation (oncogene-induced senescence, OIS) and genotoxic therapeutic agents (therapy-induced senescence, TIS) ^2^. Senescent cells undergo profound changes ranging from transcriptional, epigenetic and morphological to metabolic alterations, and exhibit a distinct secretome, the senescence-associated secretory phenotype (SASP) ^3, 4^. Senescence-associated stable cell cycle arrest underpins a number of pathophysiological conditions, but the vast majority of age-related disorders are attributable to the capacity of senescent cells to influence their microenvironment through development of the SASP ^4^.

Activation of senescent cell anti-apoptotic pathways (SCAPs), can effectively prevent these cells from undergoing programmed cell death, despite the accumulation of damage to DNA and other macromolecules ^5^. SCAPs and associated pathways activated in senescent cells present therapeutic targets for development of senotherapeutic agents, including drugs that induce selective elimination of senescent cells (senolytics) or suppress release of tissue-damaging, pro-inflammatory SASP factors (senomorphics) ^6^. Advances in understanding aging and senescence have provided a basis for developing senolytics and senomorphics, which hold the potential to intervene against various age-related pathologies and to achieve a healthy longevity ^7^. There have been challenges in developing senolytic drugs that are safe, efficacious, and have lasting effects. Efforts to translate such agents into clinical application have begun recently.

Senomorphic treatments are an alternative therapeutic approach for targeting senescent cells through limiting detrimental effects of the SASP without directly eliminating senescent cells. Early senomorphics were mainly discovered by serendipity, including metformin and rapamycin ^8^. Many natural products, particularly polyphenols, exhibit antioxidant and anti-inflammatory activities, and potentially suppress senescence-associated phenotypes by attenuating the pro-inflammatory SASP. Several naturally occurring flavonoids including apigenin and kaempferol can suppress SASP factors in bleomycin-induced senescent BJ fibroblasts ^9^. Studies focusing on synthetic flavones suggested that hydroxyl substitutions at C-2’,3’,4’,5 and 7 are critical for inhibiting the SASP ^9^.

Alzheimer’s disease (AD) is a neurodegenerative disorder associated with presence of amyloid-β (Aβ)-containing plaques and tau-containing neurofibrillary tangles ^10^. As the most prevalent cause of dementia, AD affects over 35 million people worldwide ^11^. Development of pharmacological agents for AD has been slow and expensive, with an overall rate of treatment failure greater than 99% ^12^. To date, Although there are FDA-approved anti-amyloid therapies, optimal strategies to prevent or delay the onset or slow the progression of AD remain highly desired. AD drug discovery entails target identification, followed by high-throughput screening and lead optimization of new compounds. Advanced age has been proposed as the greatest risk factor for AD, with the link between age and AD becoming increasingly clear. Cells displaying senescent features are identified in brains of AD patients and animals, implying a causal role of senescence in AD pathogenesis ^13–15^.

Dihydromyricetin (DMY) is a natural flavonoid with biological activities including potential protection against brain aging-related dysfunction by modulating oxidative stress and inflammation-related senescence of hippocampal neurons ^16^. DMY can alleviate asthma, osteoporosis, kidney injury, nephrotoxicity, atherosclerosis, myocardial infarction, depressive disorders and behavioral deficits ^17–19^. Although these effects imply a role for DMY in disease prevention and treatment, insights into key mechanisms of DMY in maintaining health and curtailing pathologies remain limited. In this study, we screened a natural medicinal agent (NMA) library and identified candidate phytochemicals including DMY as being potential novel senotherapeutic agents. We found that DMY, a flavonoid compound, has senomorphic effects in assays involving stromal cell lines such as fibroblasts and endothelial cells. In contrast, DMY exhibited a senolytic potential against microglial cells, suggesting it has pleiotropic senotherapeutic effects. DMY combined with DNA damaging chemotherapy reduced tumor sizes and improved treatment efficacy. In preclinical studies involving a 5xFAD mouse model, DMY reduced Aβ accumulation and improved cognitive ability. We propose that DMY may be a novel phytochemical senotherapeutic agent with the potential for delaying age-related dysfunction and mitigating age-related diseases.

## Results

### Drug screening reveals the potential of DMY as a senotherapeutic agent

A handful of medicinal herbs and bioactive components derived from plant sources have shown geroprotective properties and therapeutic potential ^7, 20^. To identify novel senotherapeutic compounds targeting senescent cells, we conducted unbiased drug screening of a library composed of 50 natural medicinal agents (NMAs), most of which are phytochemical compounds *per se* (Supplementary Table 1). In this study, we employed a primary normal human prostate stromal cell line, PSC27, as a cell-based model which comprises predominantly fibroblasts and exhibits fibroblast characteristics ^21^. PSC27 develops a typical pro-inflammatory SASP upon exposure to stresses such as genotoxic chemotherapy and ionizing radiation ^21, 22^. To induce cellular senescence, we treated cells with pre-optimized sub-lethal doses of either bleomycin (BLEO, 50 μg/mL) or ionizing radiation (RAD, 10 Gy), and observed increased staining positivity of senescence-associated β-galactosidase (SA-β-Gal), decreased BrdU incorporation and elevated DNA damage response (DDR) foci 8-10 days afterwards (Supplementary Fig. 1a-c). Similar effects on human umbilical vein endothelial cells (HUVECs) and human breast fibroblasts (HBF1203) were observed, including increased SA-β-Gal staining positivity (Supplementary Fig. 1d, e). Thus, these cells appear to be an effective model for studying cellular senescence.

We established an *in vitro* screening strategy to assess the effects of individual phytochemical agents on two parameters: senescent cell expression profile (at 50 μM) and senescent cell viability (at 100 μM). Procyanidin C1 (PCC1) served as a positive senolytic control (Supplementary Fig. 2a, b). Our preliminary data indicated that a number of these agents were able to downregulate the SASP when used at 50 μM (Supplementary Fig. 2c, d). However, thorough screening of the NMA library failed to identify any senolytic agents that can selectively eliminate senescent cells, as was the case for PCC1 (100 μM) (Supplementary Fig. 2b-f). Our results not only imply the scarcity of naturally available senolytics but also indicate challenges in expanding the arsenal of this specific subclass of senotherapeutics. Although no new senolytic compounds were identified, several NMA components exhibited senomorphic activity, including dihydromyricetin (DMY), warranting further investigation (Supplementary Fig. 2c, d and Supplementary Fig. 3a-c).

When DMY was used at higher concentrations such as 400 μM, we observed significantly reduced survival of PSC27 cells, no matter if cells were senescent or not, suggesting that this concentration of DMY is cytotoxic (Fig. 1a, b). Thus, DMY may not be an ideal senolytic agent for fibroblasts. However, DMY exhibited senomorphic activity at 100 μM, as evidenced by attenuated SASP factor expression (Fig. 1c, d). Despite this, it preserves other senescence features such as nuclear membrane degradation and cell cycle arrest (Supplementary Fig. 5a-e). RNA sequencing (RNA-seq) data showed that DMY comprehensively modulated gene expression in senescent cells, as it upregulated 1152 and downregulated 2044 genes, respectively, while 4059 and 1959 genes were upregulated and downregulated, respectively, during cellular senescence (Supplementary Fig. 4a, b). Heatmap, volcano plot and GSEA analyses revealed significant gene expression alterations induced by DMY, including changes in the SASP signature (Fig. 1e-h). DMY predominantly affected pathways related to DNA damage- and telomere stress-induced senescence, as well as the NF-κB pathway, with most of these changes being reversed by DMY treatment (Supplementary Fig. 4c). Gene Ontology (GO) profiling indicated that the most affected biological processes, cellular components, and molecular functions were associated with the pro-inflammatory or secretory phenotype of senescent cells, while Kyoto Encyclopedia of Genes and Genomes (KEGG) analysis suggested that the PI3K-Akt and MAPK signaling pathways were among the most affected axes by DMY (Fig. 1i).

**Fig. 1.**
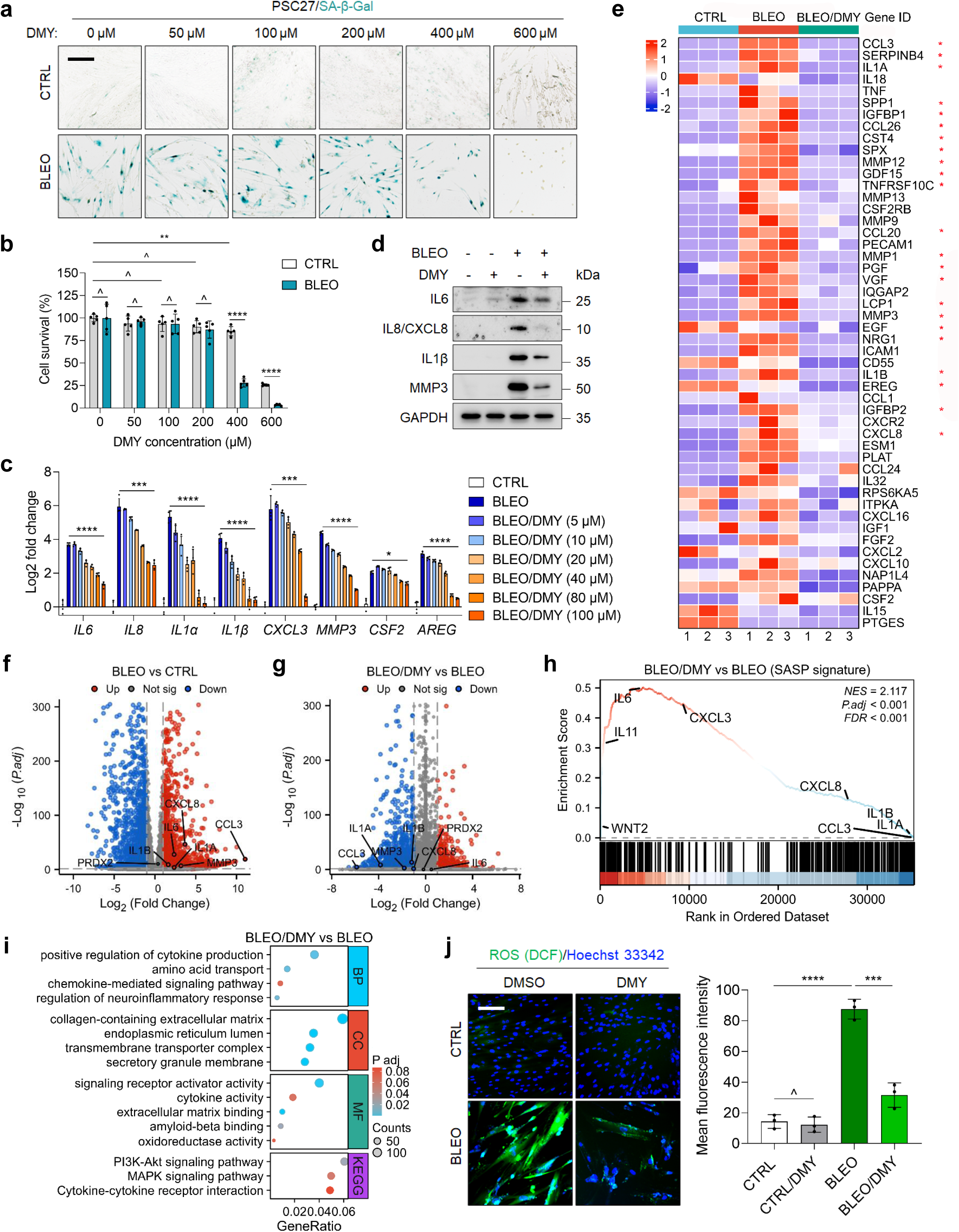
DMY exhibits senomorphic activity in fibroblasts. (**a**) Representative images of SA-β-Gal staining of proliferating (CTRL) and senescent (BLEO) PSC27 cells exposed to increasing concentrations of DMY (0-600 μM). Cellular senescence was induced by treatment with 50 μg/ml BLEO (bleomycin) for 12 h, followed by 10 d to allow cells to senesce. Scale bar, 20 μm. (**b**) Viability of CTRL and BLEO PSC27 cells following treatment by increasing concentrations DMY (0-600 μM). (**c**) Quantitative analysis of typical SASP factor expression at transcription level, comparing BLEO-induced senescence with increasing concentrations of DMY treatment (0-100 μM). (**d**) Immunoblot analysis of typical SASP factors at protein level in PSC27 cells upon BLEO-induced senescence with or without 100 μM DMY treatment. GAPDH, loading control. (**e**) Heatmap depicting the top 50 senescence/SASP-associated genes from the SenMayo geneset significantly downregulated upon treatment by 100 μM DMY. Red stars indicate representative SASP factors. (**f**) Volcano plot displaying differentially expressed genes in PSC27 cells following BLEO treatment. SASP factors are highlighted. (**g**) Volcano plot analysis of differentially expressed genes in senescent PSC27 cells after DMY treatment. SASP factors are marked. (**h**) GSEA profiling of typical soluble factors in the broad spectrum of SASP. (**i**) GO and KEGG pathway enrichment analysis was performed with 3,196 differentially expressed genes identified after DMY treatment of senescent PSC27 cells. (**j**) ROS levels were measured using DCFH-DA probe (green) in CTRL and BLEO PSC27 cells, with either DMSO or 100 μM DMY treatment for 10 d. Hoechst 33342 (blue) used for nuclear staining. Left, representative images. Right, comparative statistics. Scale bar, 20 μm. Unless otherwise stated, data are presented as mean ± SD from 3 independent biological replicates. *P* values were calculated using Student’s *t*-tests. ^, *P* > 0.05; *, *P* < 0.05; **, *P* < 0.01; ***, *P* < 0.001; ****, *P* < 0.0001.

To test further the data from BLEO-induced senescence, we examined the effect of ionizing radiation (RAD) and observed a similar pattern (Supplementary Fig. 4d). Furthermore, we compared DMY’s efficacy to established SASP inhibitors, such as the flavonoid rutin and the small molecule compound SB203580 ^23, 24^, and observed comparable results (Supplementary Fig. 4e). Next we employed HUVEC and HBF1203 cells to further assess effects of DMY and found that DMY caused concentration-dependent inhibition of the SASP (Supplementary Fig. 4f-k). The results further indicate that DMY has efficacy as a broad-spectrum senomorphic agent. Notably, we observed a significant reduction of reactive oxygen species (ROS) levels following DMY treatment (Fig. 1j), consistent with its antioxidant activity^25^. As one of the most abundant natural flavonoids in plants such as vine tea, DMY has demonstrated a wide range of pharmacological effects ^25^. In addition to the general properties of flavonoids, DMY exhibits potential anti-diabetes, anti-tumor, cardioprotection, hepatoprotection, neuroprotection and dermatoprotection activities ^26–28^. However, the geroprotective potential and functional mechanism of DMY, particularly its role in modulating senescent cells and their bioactivities including SASP regulation, remain hitherto underexplored.

### DMY exhibits senolytic activity against microglial cells

Given that DMY acts as an effective senomorphic agent for certain cell types such as fibroblasts and endothelial cells, we asked whether it exhibits distinct activities, such as a senolytic properties in other cell lineages. We noted that DMY significantly reduced SA-β-Gal positivity and the number of senescent HMC3 cells, an established human microglia cell line, starting at 50 μM (Fig. 2a, b). Immunoblots showed that DMY promoted apoptosis of HMC3 cells, as evidenced by increased cleavage of caspase 3 and elevated expression of both PMAIP1 (NOXA), BBC3 (PUMA) and BAX, key apoptosis regulators, while BCL2 levels remained unchanged (Fig. 2c). In contrast to the 3,196 significantly expression-altered genes induced by DMY in senescent PSC27 cells, RNA-seq analysis identified 634 upregulated and 461 downregulated genes in DMY-treated senescent HMC3 cells (fold change > 2, *P* < 0.05) (Supplementary Fig. 6a, b). These GSEA data suggest that HMC3 cells exhibit activities including DNA damage response and secretion during cellular senescence, resembling those of PSC27 cells (Supplementary Fig. 6c-g). However, HMC3 cells did not exhibit a typical SASP upon senescence induction (Fig. 2d). Consistent with GSEA results indicating DMY-induced programmed cell death (Fig. 2e), volcano plot analysis revealed significant upregulation of the apoptosis-regulatory genes NOXA and PUMA (Fig. 2f). We next mapped the significantly changed transcripts to the GO and KEGG databases comprising Entrez Gene, HPRD and UniProt accession identifiers ^29–31^. Analysis indicated that the most prevalent biological processes (BP) altered by DMY were the ERK1/ERK2 cascade and aging, while molecular functions (MF) were primarily related to cytokine/chemokine activity and oxidoreductase activity (Fig. 2g). Additionally, flow cytometry analysis after Annexin V-FITC/PI double staining showed that DMY significantly induced apoptosis in senescent HMC3 cells (Fig. 2h). JC-1 staining revealed that DMY decreased mitochondrial membrane potential (MMP) in senescent HMC3 cells, as evidenced by reduced J-aggregate levels (red fluorescence intensity) (Fig. 2i). Notably, in contrast to PSC27 cells, fluorescent probe DCFH-DA staining showed that DMY significantly increased ROS levels in senescent HMC3 cells (Fig. 2j). Collectively, these results indicate that DMY has senolytic activity in microglial cells, mediated largely through apoptosis induction, MMP decline and ROS bioproduction.

**Fig. 2.**
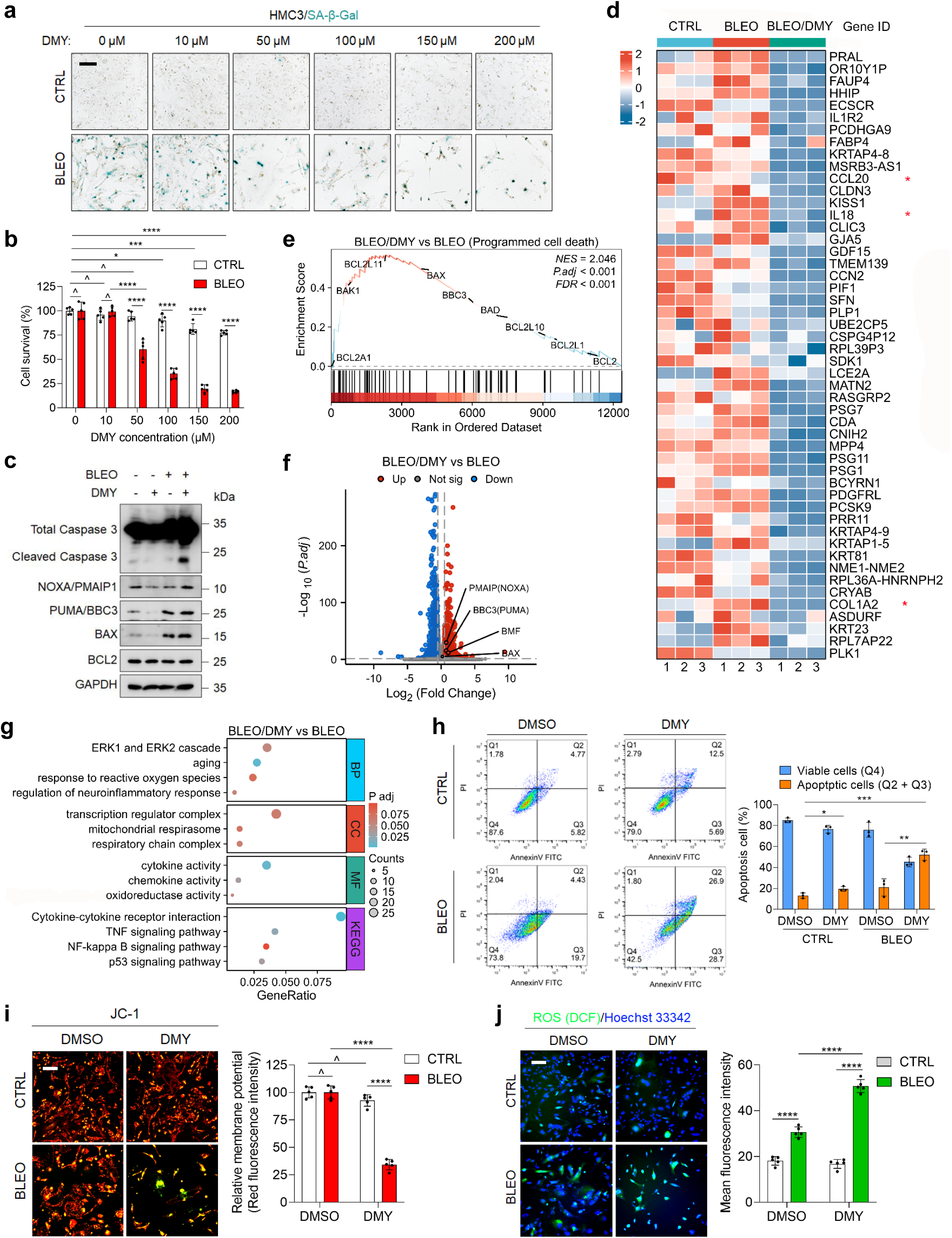
DMY shows senolytic activity against microglial cells. (**a**) Representative images of SA-β-Gal staining performed on non-senescent (CTRL) and senescent (BLEO) HMC3 cells exposed to increasing concentrations of DMY (0-200 μM). Cellular senescence was induced by 50 μg/ml BLEO treatment for 12 h, followed by 7 d to allow senescence development. Scale bar, 20 μm. (**b**) Viability of both proliferating (CTRL) and senescent (BLEO) HMC3 cells treated with increasing concentrations of DMY (0-200 μM). (**c**) Immunoblot analysis of apoptosis-related signaling molecules in HMC3 cells upon BLEO-induced senescence with or without 100 μM DMY treatment. GAPDH, loading control. (**d**) Heatmap showing the top 50 human genes significantly downregulated by 100 μM DMY treatment in HMC3 cells. Red stars, representative SASP factors. (**e**) GSEA profiling of programmed cell death in DMY-treated senescent HMC3 cells. (**f**) Volcano plot of differentially expressed genes in senescent HMC3 cells following DMY treatment, with apoptosis-associated factors highlighted. (**g**) GO and KEGG pathway enrichment analysis of 1,095 differentially expressed genes identified after DMY treatment of senescent HMC3 cells. (**h**) Apoptosis levels in CTRL and BLEO HMC3 cells treated with either DMSO or 100 μM DMY for 7 d, assessed by Annexin V-FITC/PI staining and flow cytometry. Quantification shows percentages of viable (Q4, double-negative) and apoptotic (Q2/Q3, Annexin V+/PI±) populations. Left, representative plots. Right, statistical comparison. (**i**) MMP measured by JC-1 staining in CTRL and BLEO HMC3 cells treated with either DMSO or 100 μM DMY for 7 d. Red fluorescence indicates JC-1 aggregates in intact mitochondria, while green fluorescence represents JC-1 monomers in the cytosol (early apoptosis). Left, representative images. Right, statistical comparison. Scale bar, 20 μm. (**j**) ROS levels detected by DCFH-DA staining (green) in CTRL and BLEO HMC3 cells with either DMSO or 100 μM DMY treatment for 7 d. Nuclei were counterstained with Hoechst 33342 (blue). Left, representative images. Right, quantitative comparison. Scale bar, 20 μm. Unless otherwise stated, data are presented as mean ± SD from 3 independent biological replicates. *P* values were calculated using Student’s *t*-tests. ^, *P* > 0.05; *, *P* < 0.05; **, *P* < 0.01; ***, *P* < 0.001; ****, *P* < 0.0001.

Given the potential senolytic effects of DMY on microglial cells, we extended our assays to other glial cell types, including oligodendrocytes, astrocytes and murine microglia. Notably, DMY demonstrated senolytic activity against the murine microglial line BV2 (Supplementary Fig. 6g, h), resembling the effect observed in human microglial line HMC3. Notably, senescent MO3.13 oligodendrocytes exhibited apoptotic responses starting at 100 μM DMY, but their non-senescent counterparts were similarly affected at the same concentration (Supplementary Fig. 6i, g). Similarly, senescent SVG p12 astrocytes underwent apoptosis at a DMY concentration of 200 μM, but their non-senescent counterparts responded at the same concentration (Supplementary Fig. 6k, l). Since both senescent and non-senescent cells exhibited similar concentration-dependent responses, our results suggest that DMY cannot serve as an ideal senolytic agent for oligodendrocytes or astrocytes.

### DMY directly binds to PRDX2 as a molecular target in senescent cells

Given the efficacy of DMY in suppressing SASP secretion across certain cell types upon senescence while promoting apoptosis in microglial cells upon senescence, we queried underlying mechanisms of the senomorphic and senolytic activities of DMY. To this end, we employed two methodologies: human proteome microarray and pull-down, the latter combined with liquid chromatography-mass spectrometry/mass spectrometry (LC-MS/MS) assays to identify potential intracellular factors that directly interact with DMY. First, we chose to use biotin-labeled DMY (DMY-Biotin) to assess its binding affinity to recombinant proteins coated on the surface of HuProt^TM^ 20K human protein microarrays (Fig. 3a and Supplementary Fig. 7a). We reasoned that proteins binding to DMY-Biotin may represent direct targets of DMY. After applying DMY-Biotin to the HuProt^TM^ 20K human protein microarray, we identified 395 proteins with significant specific binding (Fig. 3b, c and Supplementary Fig. 7b), the majority of which were associated with antioxidant and oxidoreducatase activity according to GO and KEGG analyses (Supplementary Fig. 7c). Additionally, we applied pull-down by incubating DMY-Biotin with lysates from two cell lines, PSC27 and HMC3, which were induced to become senescent, then performed LC-MS/MS analyses to further identify potential target proteins (Fig. 3d). Venn and volcano plot analyses identified a total of 130 potential target proteins shared between senescent PSC27 and HMC3 cells (Fig. 3e, f). Interestingly, GO and KEGG analyses revealed a distinct DMY-binding pattern in PSC27 cells, namely DMY-pull-down proteins primarily associated with DNA damage repair and nucleotide excision repair pathways (Supplementary Fig. 7d). Conversely, in HMC3 cells these proteins exhibited intensive enrichment of oxidative phosphorylation and mitochondrial functions (Supplementary Fig. 7e). Venn diagram analyses integrating data from human protein microarray alongside pull-down and LC-MS/MS techniques revealed that only 4 proteins, namely APIP, PRDX2, CHD4 and MMS19, can likely interact with DMY-Biotin (Fig. 3g). Silver staining suggested that each of these DMY-pull-down proteins has a molecular weight of approximately 52, 43, 35 and 22 KDa, respectively (Fig. 3h). We reasoned that DMY directly binds to PRDX2 (22 KDa), with the interaction subsequently confirmed by immunoblots (Supplementary Fig. 7f).

**Fig. 3.**
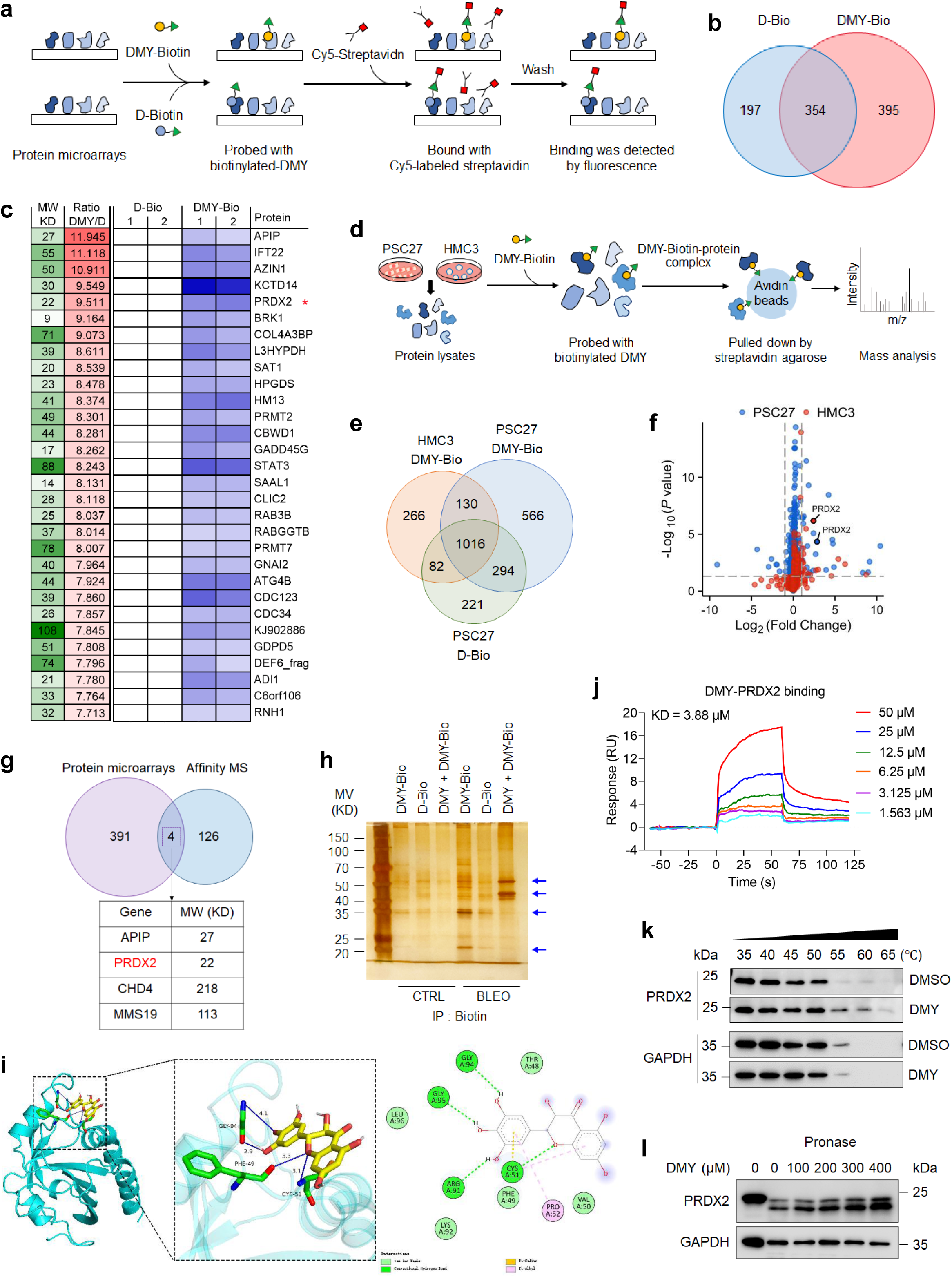
Identification of a DMY direct target PRDX2 in senescent cells. (**a**) Schematic illustration of DMY-binding protein identification with HuProt^TM^ 20K human proteome microarrays. (**b**) Venn diagram showing the 395 DMY-Biotin binding proteins identified as significant by human proteome microarrays. (**c**) Heatmap displaying the 30 human proteins with the highest binding affinity to the DMY-Biotin probe on HuProt^TM^ 20K human proteome microarrays. (**d**) Schematic illustration of the pull-down and LC-MS/MS procedures using a DMY-Biotin probe for target enrichment, followed by streptavidin-agarose pull-down and LC-MS/MS analysis to identify DMY-binding proteins in senescent PSC27 and HMC3 cells. (**e**) Venn diagram showing the 130 significant potential target proteins identified by pull-down and LC-MS/MS assays in senescent PSC27 and HMC3 cells. D-Bio was established using a biotinylated control in senescent PSC27 cells. (**f**) Volcano plot analysis of DMY-Biotin pull-down proteins in senescent PSC27 and HMC3 cells, with PRDX2 highlighted. (**g**) Venn diagram illustrating 4 potential target proteins (APIP, PRDX2, CHD4 and MMS19) identified by human proteome microarrays combined with pull-down and LC-MS/MS strategies. (**h**) Cell extracts from both non-senescent and senescent PSC27 cells were incubated with 100 μM DMY-Biotin, either alone or with 200 μM unlabeled DMY, followed by affinity capture using streptavidin-conjugated agarose beads and subsequent silver staining of the proteins eluted from agarose beads. Blue arrows indicate significantly enriched DMY-Biotin pull-down proteins. (**i**) Predicted binding mode of DMY to PRDX2 from molecular docking simulations using AutoDock. The crystallographic PRDX2 structure (PDB ID: 5IJT) served as the docking template, with Phe49 Cys51, and Gly94 highlighted in structure. (**j**) Surface plasmon resonance (SPR) demonstrated a direct interaction between DMY and PRDX2. Dissociation constants were calculated with data from two independent experiments (mean values shown). (**k**) Immunoblot showing the CETSA assay results evaluating the thermal stabilization of PRDX2 upon incubation with DMY at temperatures ranging from 35-65°C in protein lysates of PSC27 cells. GAPDH served as a negative control. (**l**) Immunoblot showing the DARTS assay results and indicating the proteolytic stability of PRDX2 at DMY concentrations 0-400 μM in 12 μg/mL pronase. GAPDH, negative control.

Despite the above findings, the interaction between PRDX2 and DMY warrants further validation. Molecular docking simulations suggested DMY has the potential to interact with the Phe49, Cys51, Arg91, Gly94 and Gly95 residues of PRDX2 (Fig. 3i). Human protein microarray disclosed that all PRDX isoforms (PRDX1-4) can interact with DMY, wherein PRDX2 produced the highest signal-to-noise ratio (SNR) (Supplementary Fig. 7g). Detailed analysis *via* surface plasmon resonance (SPR) demonstrated that recombinant human PRDX2 interacts with DMY with prominent affinity, indicated by a dissociation constant (KD) of 3.88 μM (Fig. 3j). In contrast, recombinant human PRDX1 and PRDX6 had diminished SNR in protein microarrays, yielding dissociation constants of 24.51 μM and 34.45 μM, respectively (Supplementary Fig. 7h, i). A cellular thermal shift assay (CETSA) was performed using a temperature gradient (35°C to 65°C), along with increasing concentrations of DMY at 55°C. The results indicated concentration-dependent thermal stabilization of PRDX2 upon DMY binding (Fig. 3k and Supplementary Fig. 7j). To further examine binding dynamics, we performed drug affinity responsive target stability (DARTS) with DMY (0-400 μM) in the presence of pronase, which revealed a significant concentration-dependent increase in PRDX2 stability (Fig. 3l). Collectively, our data suggest that DMY, a naturally derived flavonoid, binds to PRDX2 protein with high affinity. Thus, PRDX2 appears to be a valid molecular target of DMY.

To validate PRDX2 as the target of DMY as to its senotherapeutic properties, we employed shRNAs to downregulate PRDX2 expression in fibroblasts and microglia (Supplementary Fig. 8a, b). PRDX2 deficiency in senescent PSC27 fibroblast cells significantly decreased cell viability upon DMY treatment (Supplementary Fig. 8c, e). PRDX2 depletion in HMC3 microglial cells caused reduced cell viability even at lower DMY concentrations (Supplementary Fig. 8d, f). To the contrary, overexpression of PRDX2 in HMC3 cells abrogated DMY’s capacity in eliminating senescent microglial cells (Supplementary Fig. 8g-i). We further found reduced ROS accumulation in senescent microglial cells treated with DMY after PRDX2 overexpression (Supplementary Fig. 8j). Together, these results validate PRDX2 as a major target of DMY to exert its senolytic effect.

### DMY is senomorphic by promoting PRDX2 nuclear translocation to repair DNA damage in senescent fibroblasts

Previous studies reported that DMY activates SIRT1 in the context of diseases including AD and myocardial injury ^32, 33^. However, our immunoblot analysis revealed no significant effect on SIRT1 expression in senescent PSC27 cells. Notably, DMY significantly suppressed the ATM/p38/AKT/mTOR signaling axis, which regulates SASP production, while showing no effect on p16^INK4a^ and p21^CIP1^ expression in senescent cells (Fig. 4a). Thus, DMY effectively inhibited SASP expression of senescent cells without altering cellular senescence *per se* (Supplementary Fig. 5). Notably, immunofluorescence staining indicated that DMY significantly reduced the signal of γH2AX, a key marker of DDR, indicating its potential capacity in repairing DDR in senescent cells (Fig. 4b). Additionally, comet assay and DNA repair reporter systems further validated DMY’s capability to promote DNA repair in senescent cells (Fig. 4c, d). Thus, DMY effectively inhibited SASP expression of senescent cells without altering the cellular senescence process *per se*. Immunofluorescence staining indicated that DMY significantly reduced the signal of γH2AX, a key marker of DDR, indicating its potential to elevate DDR capacity in senescent cells (Fig. 4b). To elucidate the PRDX2-mediated biological effects of DMY, we chose to first assess the expression level of PRDX2 during cellular senescence. Previous studies reported that PRDX2, an antioxidant peroxidase, is upregulated in senescent cells^34^. To the contrary, we found that upon senescence, expression of PRDX2 and its family members (PRDX1-6) exhibited no significant changes at either the transcriptional or protein level (Supplementary Fig. 7a, b). Although DMY significantly inhibited expression of the SASP in senescent cells, it did not generate a notable impact on PRDX2 expression (Supplementary Fig. 7c, d).

**Fig. 4.**
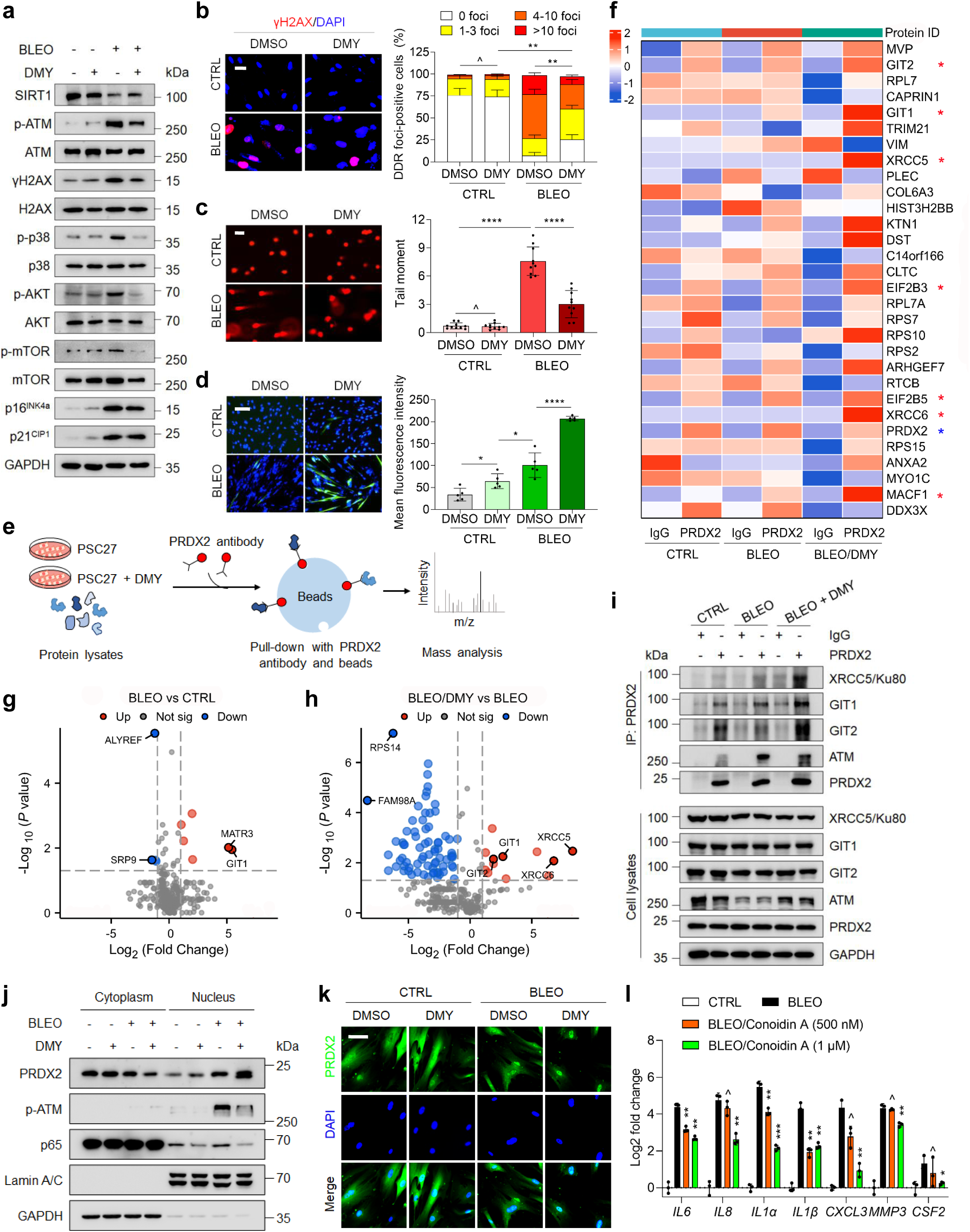
DMY promotes nuclear translocation of PRDX2 to repair DNA damage in senescent fibroblasts. (**a**) Immunoblot analysis of key factors involved in cellular senescence and SASP development in PSC27 cells cultured with or without 100 μM DMY for 10 d. GAPDH, loading control. (**b**) Left, Representative images of γH2AX immunofluorescence (IF) staining (red) in PSC27 cells cultured with either DMSO or 100 μM DMY for 10 d. Nuclei were stained with DAPI (blue). Scale bar, 10 μm. Right, Quantification of γH2AX IF intensity. Data were analyzed using two-way ANOVA. (**c**) Left, Representative images of comet assay in PSC27 cells cultured with either DMSO or 100 μM DMY for 7 d. Scale bar, 10 μm. Right, Quantification of tail moment. (**d**) Left, Representative images of DDR reporter in PSC27 cells transfected with reporter plasmid pLCN DSB, then treated with DMSO or 100 μM DMY for 7 d. Scale bar, 20 μm. Right, fluorescence intensity. (**e**) Schematic of a Co-IP assay coupled with LC-MS/MS to identify PRDX2-interacting proteins in senescent PSC27 cells. Cell lysates were immunoprecipitated with either control IgG or anti-PRDX2 antibody, followed by agarose bead pull-down and LC-MS/MS analysis. (**f**) Heatmap displaying the top 30 human proteins enriched by PRDX2 immunoprecipitation in (**e**). Red stars indicate nuclear function-associated proteins. (**g**-**h**) Volcano plots of proteins enriched by anti-PRDX2 antibody, highlighting significantly enriched proteins. (**i**) Co-IP followed by immunoblotting to validate interactions identified in (**f**). Lysates were immunoprecipitated with control IgG or anti-PRDX2, then probed for Ku80 (XRCC5), GIT1, GIT2 or ATM in both immunoprecipitates and input lysates. GAPDH, loading control. (**j**) Nuclear-cytoplasmic fractionation assay followed by immunoblotting of PRDX2, p65 and p-ATM subcellular localization in PSC27 cells treated with 100 μM DMY for 10 d. Lamin A/C and GAPDH, nuclear and cytoplasmic loading controls, respectively. (**k**) Representative immunofluorescence images of PRDX2 (green) in PSC27 cells treated with BLEO and/or 100 μM DMY for 10 d. Nuclei were stained with DAPI (blue). Scale bar, 10 μm. (**l**) Transcript analysis of PRDX2 and SASP factors in senescent PSC27 cells treated with the PRDX2 inhibitor Conoidin A for 10 days. Data represent mean ± SD from 3 independent biological replicates. *P* values were calculated using Student’s *t*-tests. ^, *P* > 0.05; *, *P* < 0.05; **, *P* < 0.01; ***, *P* < 0.001.

Subsequently, we performed PRDX2-based co-immunoprecipitation (Co-IP) in DMY-treated senescent PSC27 cells to identify PRDX2-interacting proteins by mass spectrometry (Fig. 4e). The heatmap analysis showed that proteins with significant binding affinity to PRDX2 were predominantly nuclear-localized in senescent cells (Fig. 4f). Volcano plot analysis revealed that PRDX2-interacting proteins included XRCC5, XRCC6, GIT1 and GIT2, which have been implicated in nuclear DNA damage repair ^35–37^ (Fig. 4g, h), are basically in line with our observation that DMY may facilitate DDR in senescent cells. GO analysis indicated that proteins interacting with PRDX2 are functionally associated with nucleic acid binding, while KEGG analysis further identified non-homologous end-joining as the major pathway regulated by these proteins in senescent cells (Supplementary Fig. 9e). Co-IP and immunoblot assays further confirmed interactions of PRDX2 with XRCC5, GIT1, GIT2 and ATM in DMY-treated senescent cells (Fig. 4i and Supplementary Fig. 9f). Additionally, nuclear-cytoplasmic fractionation followed by immunoblotting indicated that DMY induced PRDX2 nuclear translocation and decreased the nuclear level of p65 in senescent cells (Fig. 4j), suggesting that PRDX2 enters nuclei to facilitate the DDR, while p65 exits the nucleus and suppresses the SASP. Immunofluorescence staining of PRDX2 confirmed basal nuclear PRDX2 localization during cellular senescence, which was markedly intensified after DMY treatment (Fig. 4k and Supplementary Fig. 9g). Furthermore, qRT-PCR analysis indicated that Conoidin A, a PRDX1/2 inhibitor, significantly inhibited the SASP, with effects being similar to those of DMY treatment in senescent PSC27 cells (Fig. 4l and Supplementary Fig. 9h, i). Together, these findings indicated that binding of DMY to PRDX2 promotes its nuclear translocation and interaction with DNA repair proteins, thereby enhancing DNA repair efficacy in senescent cells.

### DMY is senolytic by inhibiting PRDX2 oxidoreductase activity to induce ROS-mediated apoptosis in senescent microglial cells

We next queried the molecular mechanism underlying the senolytic activity of DMY in HMC3 cells. Similar with PSC27 cells, DMY treatment did not affect PRDX2 expression in senescent HMC3 cells (Supplementary Fig. 10a). Notably, HMC3 cells exhibited a significantly lower basal level of PRDX2 than other cell types, including fibroblasts and endothelial cells (Supplementary Fig. 10b). Previous studies reported that inhibition of the oxidoreductase activity of PRDX2 can induce cancer cell apoptosis ^38^. Molecular docking simulations suggested an interaction of DMY with Cys51 of PRDX2 (Fig. 3i). This residue, together with Cys172, forms the enzyme’s catalytic site with oxidoreductase activity ^39^. As an oxidoreductase, PRDX2 catalyzes H_2_O_2_ reduction to water in the presence of NADPH ^38^. To verify the inhibitory effect of DMY on PRDX2 enzymatic activity, we applied an Amplex Red coupled spectrophotometric method as described previously ^38^. Our results indicated that enhancing the concentration of PRDX2 reduced H_2_O_2_ levels (Supplementary Fig. 10c). Importantly, DMY was found to inhibit the activity of recombinant human PRDX2 (Fig. 5a), mimicking the effect of Conoidin A, a PRDX2 inhibitor (Fig. 5b).

**Fig. 5.**
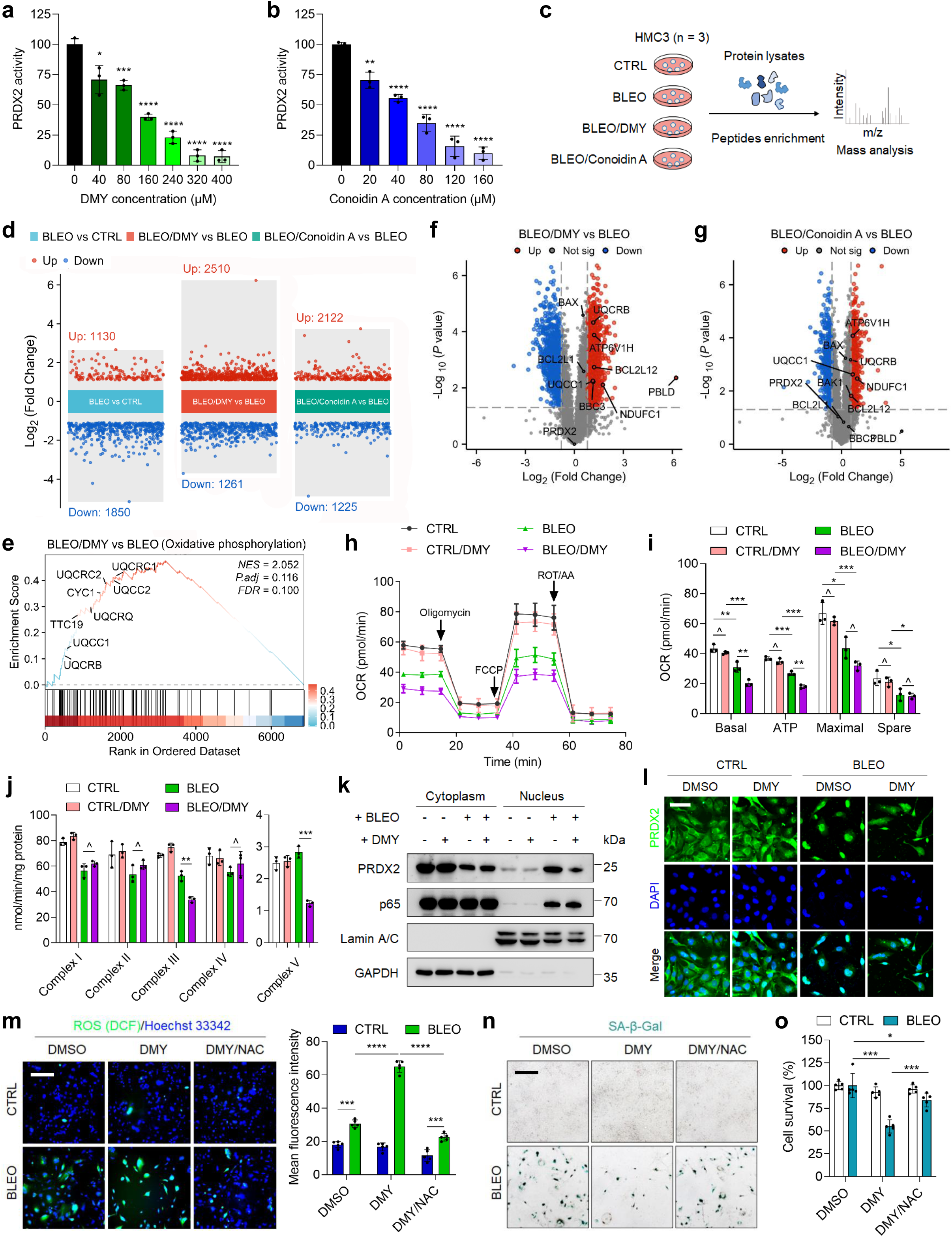
DMY induces ROS-mediated apoptosis by inhibiting PRDX2’s oxidoreductase activity in senescent microglial cells. (**a**-**b**) Recombinant human PRDX2 proteins were incubated with increasing concentrations of DMY (**a**) or Conoidin A (**b**) for 5 min, with PRDX2’s oxidoreductase activity measured. Conoidin, positive control. (**c**) Schematic of quantitative proteomics in HMC3 cells. Cells were treated with 50 μg/ml BLEO for 12 hr to induce senescence, followed by 100 μM DMY or 4 μM Conoidin A treatment for 7 d, before lysed for LC-MS/MS analysis of enriched peptides. (**d**) Multi-group volcano plots display differentially expressed proteins across treatment conditions, with each subplot representing a pairwise comparison. (**e**) GSEA revealed oxidative phosphorylation associated geneset enrichment in DMY-treated senescent HMC3 cells. (**f**-**g**) Volcano plots of proteins identified in senescent HMC3 cells treated with 100 μM DMY (**f**) or 4 μM Conoidin A (**g**), highlighting ROS production-associated proteins. (**h**) Seahorse assay analysis of oxygen consumption rate (OCR) in non-senescent (CTRL) and senescent (BLEO) HMC3 cells treated with or without 100 μM DMY for 7 d. (**i**) Quantification of basal respiration, ATP production, maximal respiration and spare respiratory capacity in (**h**). (**j**) Measurement of mitochondrial complex I-V activities in CTRL and BLEO HMC3 cells treated with or without 100 μM DMY for 5 d. (**k**) Nuclear-cytoplasmic fractionation and immunoblot analysis of PRDX2 and p65 subcellular localization in HMC3 cells treated with 100 μM DMY for 7 d. Lamin A/C and GAPDH served as nuclear and cytoplasmic loading controls, respectively. (**l**) Representative immunofluorescence images of PRDX2 (green) in HMC3 cells treated with BLEO and/or 100 μM DMY for 7 d. Nuclei were stained with DAPI (blue). Scale bar, 20 μm. (**m**) ROS levels detected by DCFH-DA staining (green) in CTRL and BLEO HMC3 cells treated with DMSO, 100 μM DMY or 100 μM DMY/5 mM NAC for 7 d. Left, representative images. Right, quantification. Scale bar, 50 μm. (**n**) SA-β-Gal staining in CTRL and BLEO HMC3 cells treated as described in **m**. (**o**) Cell viability assessed by CCK-8 assay in HMC3 cells treated as described in **m**. Data are presented as mean ± SD from 3 independent biological replicates. *P* values were calculated using Student’s *t*-tests. *, *P* < 0.05; **, *P* < 0.01; ***, *P* < 0.001; ****, *P* < 0.0001.

We found that Conoidin A exhibits senolytic activity, as evidenced by significantly reduced in cell viability of BLEO-induced senescent HMC3 cells (Supplementary Fig. 10d, e). To elucidate the molecular mechanism by which DMY induces HMC3 cell apoptosis through inhibition of PRDX2’s oxidoreductase activity, we conducted quantitative proteomics to compare the effects of DMY and Conoidin A (Fig. 5c). Quantitative proteomics revealed that DMY modified the expression of 3,771 proteins (2,510 upregulated, 1,261 downregulated), while Conoidin A modified 3,347 changes (2,122 upregulated, 1,225 downregulated) in senescent HMC3 cells (Fig. 5d). Venn analysis showed 2,886 common protein alterations, comprising 76.5% (DMY) and 86.2% (Conoidin A) of differentially expressed proteins (Supplementary Fig. 10f). Hierarchical clustering heatmap and principal component analysis (PCA) indicated nearly identical protein expression patterns between DMY- and Conoidin A-treated cells, suggesting that DMY exerts pharmacological effects in a way similar to Conoidin A in senescent HMC3 cells (Supplementary Fig. 10g, h). GSEA analysis identified enriched pathways in DMY-treated senescent HMC3 cells, including oxidative phosphorylation, mitochondrial complex I/III assembly, and ROS/RNS production (Fig. 5e and Supplementary Fig. 10i). GO analyses indicated that the most significantly affected biological processes (BP) were apoptotic signaling, mitochondrial respiratory chain complex assembly, and apoptotic mitochondrial changes, while KEGG analysis highlighted reactive oxygen species (ROS), oxidative phosphorylation, apoptosis, and p53 signaling as major pathways altered by DMY (Supplementary Fig. 10j). Volcano plots revealed upregulation of ROS-associated proteins, including UQCC1, UQCRB, NDUFC1, and ATP6V1H, following DMY and Conoidin A treatment in senescent HMC3 cells (Fig. 5f, g). Since the mitochondrial electron transport chain (ETC) is a primary source of ROS ^40^, we assessed mitochondrial function *via* oxygen consumption rate (OCR) measurement through seahorse assays. Senescent HMC3 cells displayed diminished basal respiration and ATP production as compared with non-senescent cells. Treatment with DMY further decreased basal respiration and ATP production in senescent HMC3 cells, indicating DMY-induced mitochondrial dysfunction (Fig. 5h, i). Moreover, DMY was observed to reduce the activity of mitochondrial complex III and complex V, indicating compromised ETC function (Fig. 5j). Taken together, our data suggest that DMY interacts with PRDX2, inhibiting its oxidoreductase activity, disrupting redox homeostasis and impairing ETC function. These changes ultimately resulted in elevated ROS-mediated apoptosis in senescent HMC3 cells.

Subsequently, nuclear-cytoplasmic fractionation followed by immunoblotting and immunofluorescence staining showed that unlike in PSC27 cells, PRDX2 significantly localized to nuclei in senescent HMC3 cells. Moreover, DMY treatment retained PRDX2 in the cytoplasm in a concentration-dependent manner, suggesting that PRDX2 activity blockade inhibits PRDX2’s nuclear translocation (Fig. 5k, l and Supplementary Fig. 10k). Since DMY increased ROS levels in senescent cells, we next employed a ROS scavenger N-acetyl-cysteine (NAC) to determine whether ROS mediated DMY’s effects. DCFH-DA-based fluorescence staining confirmed that NAC reversed the increased ROS levels in DMY-treated senescent HMC3 cells (Fig. 5m). JC-1 staining indicated that NAC restored mitochondrial membrane potential (MMP) in DMY-treated senescent HMC3 cells (Supplementary Fig. 10l). SA-β-Gal staining and CCK8 assays suggested that NAC restored cell viability in DMY-treated senescent HMC3 cells (Fig. 5n, o). Collectively, these results indicate that DMY induces apoptosis in senescent HMC3 cells *via* a ROS-dependent mechanism, which is subject to reversal by ROS scavenging.

### DMY administration mitigates tissue aging and age-related physical dysfunction in prematurely aged mice

Senescent cells, along with their SASP, can be found in various tissues and organs across a range of pathophysiological states, most of which are implicated in aging changes and age-associated decline of physical health. To further examine the impact of DMY on senescent cells in organismal aging *in vivo*, we challenged mice with whole body irradiation. Three-month-old wild type (WT) mice were subjected to WBI at a sublethal dose of 5 Gy to induce substantial senescence within tissues. Following this, the mice received either a placebo or DMY treatment (20 mg/kg *via* intraperitoneal injection) twice a week for 3 months, and senescence biomarkers and physical function were subsequently tested (Fig. 6a). Mice that underwent WBI displayed physical changes, including significant greying of the fur. However, this condition was attenuated by DMY administration (Supplementary Fig. 11a).

**Fig. 6.**
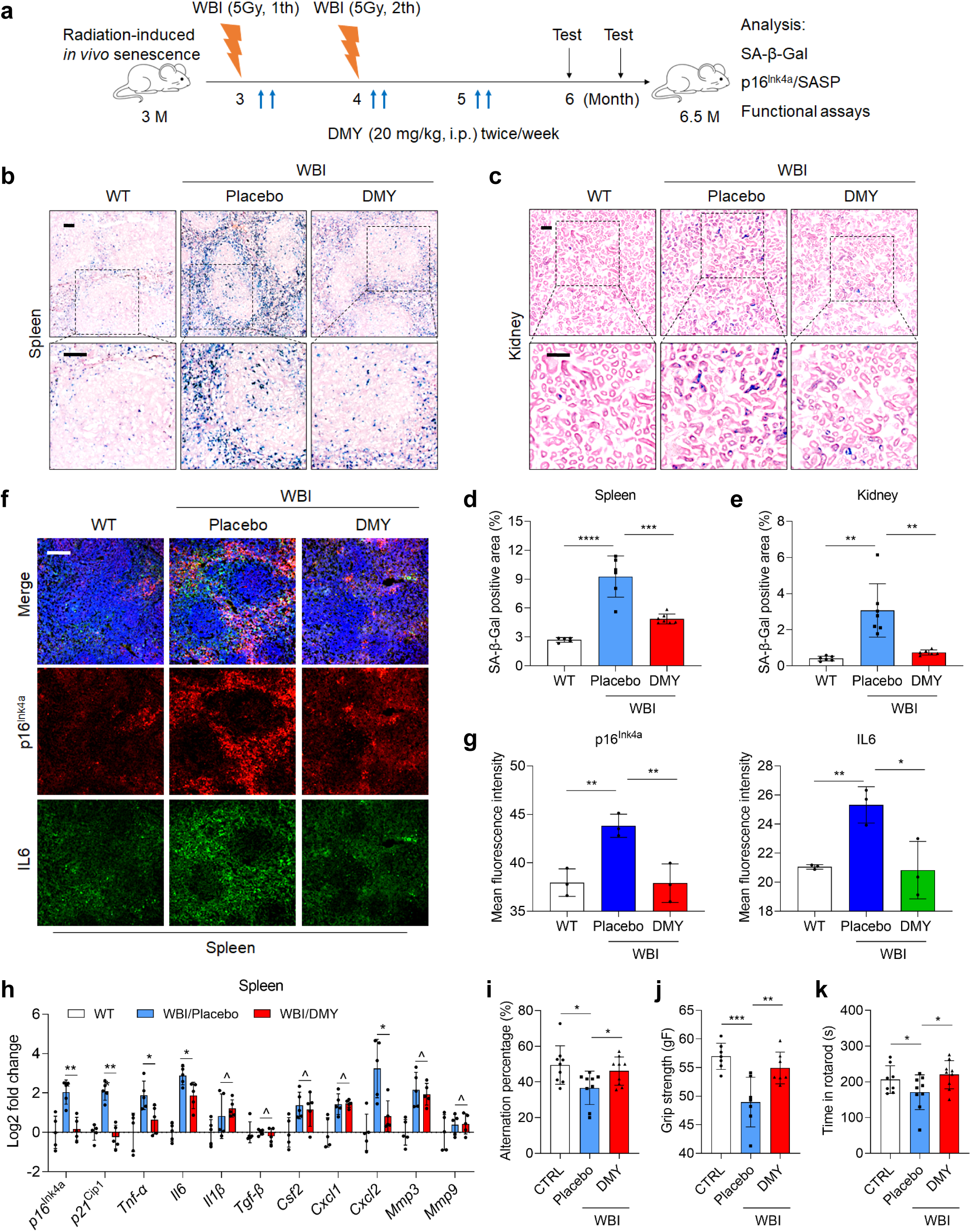
DMY administration alleviates tissue aging and physical dysfunction in prematurely aged mice. (**a**) Experimental schematic showing treatment protocol for C57BL/6J mice receiving twice 5 Gy whole body irradiation (WBI) followed by treatment with placebo or 20 mg/kg DMY twice *per* week for 3 months. (**b**-**c**) Representative SA-β-Gal staining images of spleen (**b**) and kidney (**c**) from placebo-or DMY-treated mice. Scale bar, 100 μm. (**d**-**e**) Quantitative analysis of SA-β-Gal staining intensity in spleen (**d**) and kidney (**e)** tissues, as shown in (**b**-**c**). (**f**) Representative immunofluorescence staining images of p16^Ink4a^ (red) and IL6 (green) in spleen tissues from untreated and WBI-treated mice receiving either placebo or 20 mg/kg DMY. Nuclei were stained with DAPI (blue). Scale bar, 100 μm. (**g**) Quantification of p16^Ink4a^ and IL6 immunofluorescence staining intensity in spleen tissues depicted in (**f**). (h) Quantitative analysis of transcript expression of senescence biomarkers p16^Ink4a^ and p21^Cip1^ and several representative SASP factors in spleen tissues after 3 months of placebo or 20 mg/kg DMY treatment. For each gene, signals were normalized to samples of the WT group. (**i**) Y-maze test results showing spontaneous alternation percentages in WT (n = 8), placebo (n = 9), and DMY-treated (n = 9) groups. (**j**-**k**) Physical performance assessments including grip strength (**j**) and Rotarod endurance (**k**). Data points represent individual mice. n indicates number. WBI, whole body irradiation. Data are presented as mean ± SD. *P* values were calculated using Student’s *t*-tests. ^, *P* > 0.05; *, *P* < 0.05; **, *P* < 0.01; ***, *P* < 0.001; ****, *P* < 0.0001.

In WBI-induced prematurely aged mice, there were considerably accumulated senescent cells in various tissues, as reflected by a significant increase in SA-β-Gal positivity within spleen tissues subjected to WBI (Fig. 6b, d). Notably, the data indicated that treatment with DMY led to a substantial reduction of the percentage of SA-β-Gal-positive cells, while concomitantly enhancing the structural integrity of the spleen tissues in prematurely aged mice when compared to the placebo group (Fig. 6b, d). Further examination uncovered the presence of senescent cells in several vital organs, as evidenced by elevated SA-β-Gal positivity in the kidney, liver, and pancreas tissues of these mice following WBI exposure. In contrast to the placebo-treated mice, a significant decrease in the percentage of SA-β-Gal-positive cells was observed in the DMY-treated mice (Fig. 6c, e and Supplementary Fig. 11b, c). Immunofluorescence staining provided compelling evidence that DMY treatment markedly reduced the expression of senescence markers, such as p16^Ink4a^, and canonical SASP markers such as IL6, in the spleen tissues of WBI-induced aged mice when compared to the placebo (Fig. 6f, g).

Consistent with immunofluorescence staining results in spleen tissues, DMY treatment in WBI-induced aged mice resulted in a significantly downregulated expression of senescence markers, including CDK inhibitors p16^Ink4a^ and p21^Cip1^, as well as some but not all proinflammatory factors, as determined by RT-qPCR analysis (Fig. 6h). Unlike the spleen tissues, we observed no substantial alteration in the immunofluorescence staining for p16^Ink4a^ and IL6 in kidney tissues of WBI-induced aged mice in the DMY treatment group as compared with the placebo, suggesting a heterogeneity of senescent cells (Supplementary Fig. 11d, e). Nevertheless, the transcript expression levels of the senescence biomarker p16^Ink4a^ and a variety of proinflammatory cytokines were significantly reduced in the kidney, liver and pancreas tissues of WBI-induced aged mice following DMY treatment compared to the placebo group (Supplementary Fig. 11f-h). Collectively, these results indicated that DMY reduced the accumulation of senescent cells and downregulated the expression of senescence markers in WBI-induced prematurely aged mice.

Next, we evaluated the potential of DMY administration to enhance the physical functions of WBI-induced prematurely aged mice. In comparison to aged mice that received placebo treatment, animals treated with DMY demonstrated marked recovery of short-term memory, as indicated by results from Y-maze assessments (Fig. 6i). Furthermore, DMY-treated aged mice showed a notable increase in grip strength (Fig. 6j) and exhibited superior performance in Rotarod test (Fig. 6k). These findings suggest that DMY administration is an effective intervention to mitigate physical impairments and avert cognitive decline associated with premature aging, ultimately enhancing overall health condition.

### DMY combined with chemotherapy promotes tumor regression

Given the efficacy of DMY in restraining SASP expression across certain cell types upon senescence while promoting apoptosis in senescent microglial cells, we queried its effects on tumor progression. Former studies demonstrated that senescent stromal cells secrete multiple soluble factors or metabolites to support tumor progression ^22, 24, 41, 42^. We were promoted to determine whether DMY treatment of stromal cells can affect the potential of their conditioned medium (CM) in altering malignant phenotypes of cancer cells. To this end, PSC27 stromal cells were treated with BLEO to induce senescence, with or without DMY. After a 3-d incubation, CM derived from PSC27 cells were collected and used to treat multiple prostate cancer (PCa) cell lines, including PC3, DU145, M12 and LNCaP (Supplementary Fig. 12a). Consistent with previous findings, exposure to senescent PSC27 cell-derived CM substantially increased the proliferative capacity of PCa cells (Supplementary Fig. 12b). The proliferative advantage was accompanied by enhanced migratory and invasive activities (Supplementary Fig. 12c, d). However, DMY pretreatment of stomal cells significantly reduced these pro-tumorigenic effects of stromal cell-derived CM (Supplementary Fig. 12b-d).

Since DMY downregulates SASP expression in senescent stromal cells and inhibits cancer cell malignancy *in vitro*, we sought to investigate its potential therapeutic effects on tumors *in vivo*. To simulate clinical settings, we established xenograft tumors by co-injecting PSC27 sublines with PC3 cells at an optimized ratio of 1:4 into NOD/SCID mice through tissue recombination. After confirming successful engraftment 2 weeks post-implantation, intervention regimen was initiated with weekly administrations of placebo, MIT and/or DMY starting at weeks 3, 5 and 7, continuing through an 8-week therapeutic period. Tumor sizes were measured at the end of the 8th week (Fig. 7a). Although DMY monotherapy showed modest antitumor effects, MIT treatment induced significant tumor regression, validating its chemotherapeutic efficacy. Importantly, combination therapy with DMY and MIT resulted in enhanced tumor suppression compared to MIT alone (Fig. 7b). To assess senescence induction in the tumor microenvironment, we analyzed xenograft tissues from PC3/PSC27 recombinants. Stromal cells in MIT-treated tumors displayed significant upregulation of multiple SASP components (IL6, IL8, IL1α, MMP3, GM-CSF, IL-1β, ANGPTL4 and MMP1) along with increased expression of senescence markers p16^INK4a^ and p21^CIP1^, confirming chemotherapy-induced senescence *in vivo* (Fig. 7c, d and Supplementary Fig. 13a). Histopathological examination revealed a substantial increase of SA-β-Gal-positivity following MIT treatment (Fig. 7e). Notably, while DMY co-administration did not affect the chemotherapy-induced senescence burden (Fig. 7e), it effectively attenuated SASP factor expression, as demonstrated by transcriptomic analysis (Supplementary Fig. 13b). These findings suggest that DMY specifically targets the SASP without interfering with the senescence program *per se*.

**Fig. 7.**
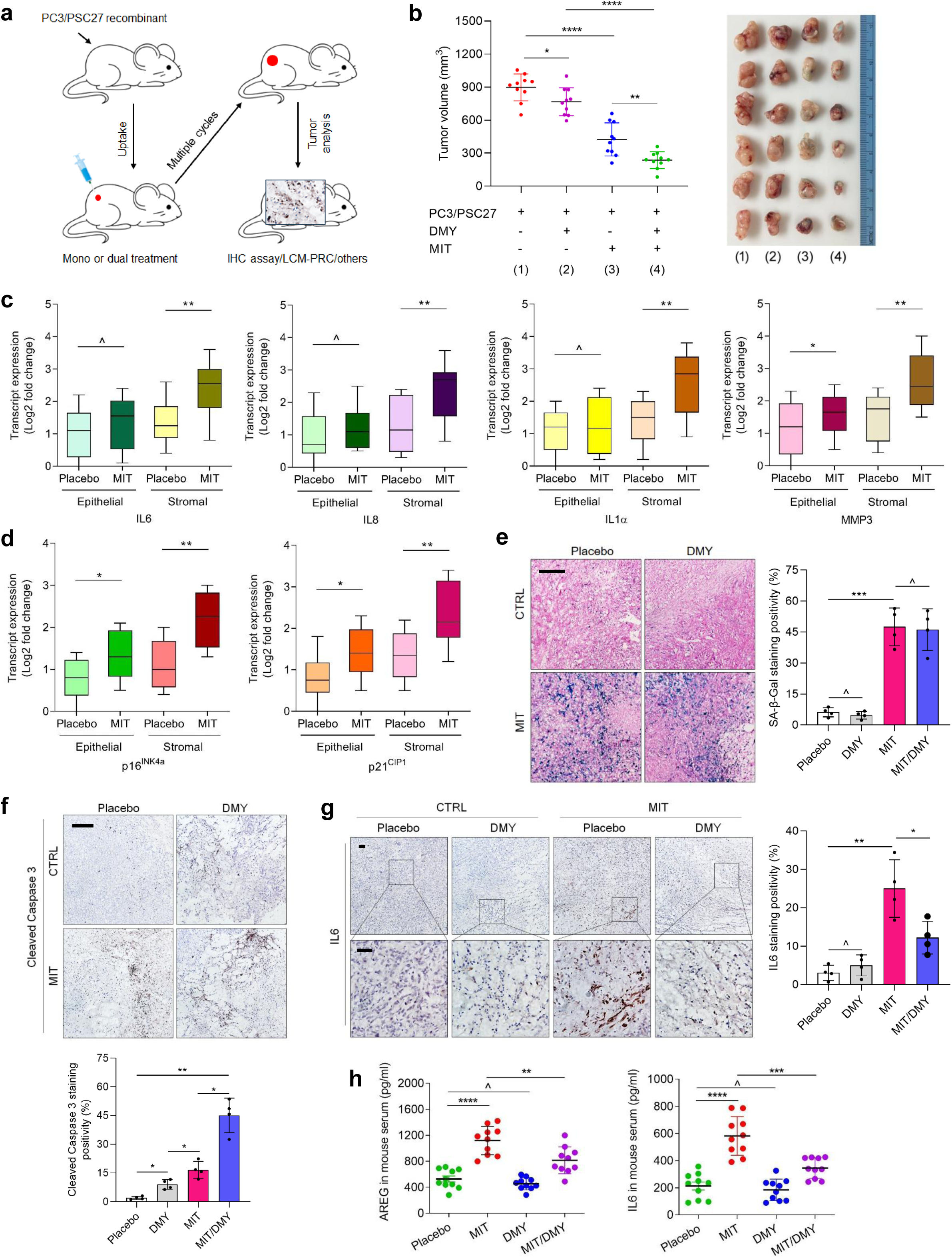
DMY combined with chemotherapy promotes tumor regression. (**a**) Schematic of a preclinical treatment procedure. Two weeks after subcutaneous inoculation and *in vivo* uptake of PC3/PSC27 cell recombinants, NOD/SCID mice received either single-agent or combinatorial metronomic treatment in multiple cycles. (**b**) Tumor end volume analysis. PC3 cells were inoculated alone or with PSC27 cells for subcutaneous implantation at mouse hind flanks, followed by 0.2 mg/kg mitoxantrone (MIT) and 20 mg/kg DMY treatment (single or combined). Left, comparative statistics. Right, representative tumor images. (**c**) Transcriptional analysis of typical SASP factors in stromal and cancer cells isolated *via* laser capture microdissection (LCM). Signals were normalized to the lowest value of placebo group. (**d**) Transcriptional analysis of two canonical senescence biomarkers p16^Ink4a^ and p21^Cip1^. (**e**) SA-β-Gal staining in tumor tissue at the end of therapeutic regimens. Left, representative images. Right, comparative statistics. Scale bar, 100 μm. (**f**) Apoptosis assessment *via* cleaved caspase 3-based immunohistochemistry staining of tumor tissues at the end of therapeutic regimen. Top, representative images. bottom, comparative statistics. Scale bar, 100 μm. (**g**) lmmunohistochemical staining of IL6 in tumor tissues at the end of therapeutic regimen. Left, representative images. Right, comparative statistics. Scale bar, 50 μm. (**i**) Circulating levels of SASP factors including AREG and IL6 in the serum of MIT/DMY-treated NOD/SCID mice. Data are presented as mean ± SD. *P* values were calculated using Student’s *t*-tests. ^, *P* > 0.05; *, *P* < 0.05; **, *P* < 0.01; ***, *P* < 0.001; ****, *P* < 0.0001.

Subsequently, we investigated the molecular mechanisms through which DMY combined with MIT contributes to tumor regression. To elucidate these pathways, we performed immunohistochemical analysis of cleaved caspase-3, an established apoptotic marker, in tumor xenograft models. Our data demonstrated that while DMY monotherapy exhibited limited cytotoxic activity, its combination with MIT substantially potentiated apoptotic cells (Fig. 7f). Further immunohistochemical examination revealed that DMY effectively counteracted the tumor-promoting effects of chemotherapy-induced SASP, as evidenced by reduced expression of IL6 and IL1β (Fig. 7g and Supplementary Fig. 13c, d). Quantitative analysis of serum SASP components through ELISA showed that MIT treatment markedly increased circulating levels of AREG and IL6, which were significantly attenuated by DMY co-administration (Fig. 7h). These results provide compelling evidence that DMY may serve as a valuable agent in combinatorial anticancer regimens, offering potential clinical benefits by modulating therapy-induced senescence. The observed pharmacological effects underscore the translational potential of DMY in improving therapeutic outcomes.

### DMY administration selectively eliminates senescent microglial cells from Aβ plaques and prevents cognitive decline in AD mice

Given the effectiveness of DMY in eliminating senescent HMC3 cell lines *in vitro*, it is tempting to determine whether DMY administration would eliminate Aβ plaque-associated senescent microglia and ameliorate cognitive dysfunction in 5×FAD mice, a transgenic model of AD. We designed an experimental protocol in which either placebo or 50 mg/kg DMY was administered once *per* day for 7 days to 12-month-old 5×FAD mice, before Aβ load, senescence biomarker expression and senescent microglia were subsequently determined (Fig. 8a). Our data indicated that this short-term DMY treatment had no significant impact on Aβ load. Instead, it significantly reduced the number of microglial cells associated with Aβ plaques, as well as the level of p16^Ink4a^ (Fig. 8b, d). Immunofluorescence staining further demonstrated that acute DMY administration triggered cellular apoptosis, as evidenced by a significant increase in cleaved caspase 3 associated with Aβ plaques (Fig. 8c, d). We noticed that there was no significant alteration in the expression of GFAP and NeuN in 5×FAD mice treated with DMY compared with the placebo group (Fig. 8d and Supplementary Fig. 14a, b), indicating that DMY treatment had no significant effect on the survival of astrocytes and neurons associated with Aβ plaques. Taken together, the data implied a target-specific effect of acute DMY treatment on senescent microglial cells associated with Aβ plaques, while largely sparing astrocytes and neurons.

**Fig. 8.**
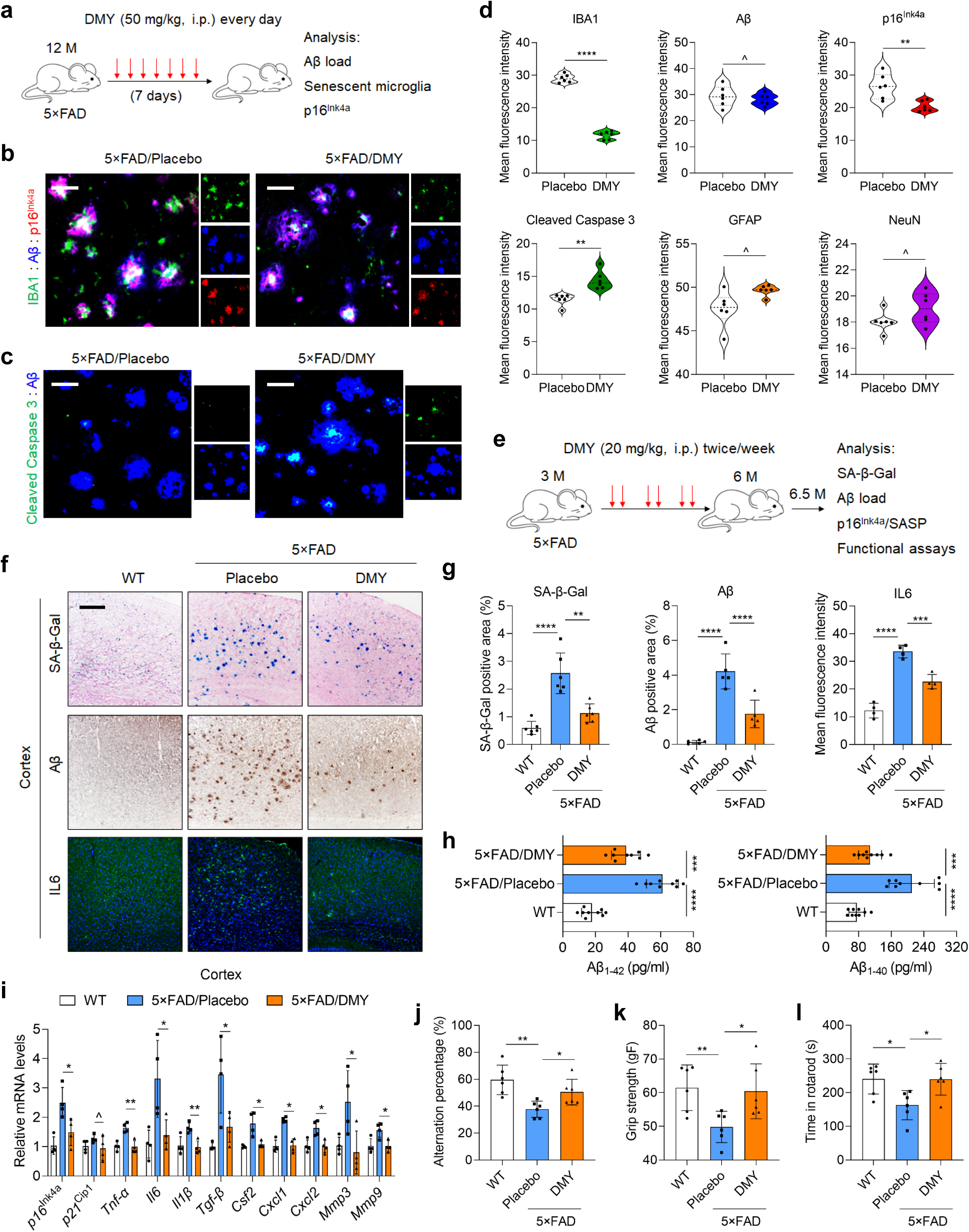
DMY administration selectively eliminates senescent microglial cells from Aβ plaques and prevents cognitive decline. (**a**) Experimental design for 7-day administration of 50 mg/kg DMY once daily in 12-month-old female 5×FAD mice. (**b**) Representative images of immunofluorescence staining for Iba1 (green), Aβ (blue) and p16^Ink4a^ (red) in brain sections of 5×FAD mice treated with either placebo or 50 mg/kg DMY for 7 d. Scale bar, 50 μm. (**c**) Representative images of immunofluorescence staining for cleaved caspase 3 (green), Aβ (blue), and p16^Ink4a^ (red) in brain sections of 5×FAD mice treated with either placebo or 50 mg/kg DMY for 7 days. Scale bar, 50 μm. (**d**) Quantification of Aβ load and immunofluorescence staining intensity of IBA1, p16^Ink4a^, cleaved caspase 3, GFAP and NeuN. (**e**) Schematic showing 3-month intermittent administration (twice weekly) of 20 mg/kg DMY or placebo in female 5×FAD mice, beginning at 3 months of age. (**f**) Representative images from cortex tissues of WT and 5×FAD mice, treated with 20 mg/kg DMY or placebo for 3 months, showing SA-β-Gal staining (blue, top), Aβ immunohistochemical staining (brown, middle) and IL6 immunofluorescence staining (green, bottom). Scale bar, 200 μm. (**g**) Quantification of SA-β-Gal staining (n = 6), Aβ plaque deposition (n = 5) and IL6 fluorescence intensity (n = 4) in cortex tissues depicted in (**f**). (**h**) ELISA-based measurement of Aβ_1-42_ and Aβ_1-40_ levels in the serum from female 5×FAD mice (n = 9) administrated with placebo or 20 mg/kg DMY. (**i**) Quantitative analysis of transcript expression of senescence biomarkers (p16^Ink4a^ and p21^Cip1^) and representative SASP factors in cortex tissues after 3-month administration with 20 mg/kg placebo or DMY. Signals were normalized to WT samples per gene. (**j**) Y-maze test showing spontaneous alternation percentages in WT (n = 6), placebo (n = 6) and DMY-treated (n = 6) groups. (**k**-**l**) Physical performance assessments including grip strength (**j**) and Rotarod endurance (**k**) (n = 6). Data points represent individual mice, while n indicates number. Data are shown as mean ± SD. *P* values were calculated using Student’s *t*-tests. ^, *P* > 0.05; *, *P* < 0.05; **, *P* < 0.01; ***, *P* < 0.001; ****, *P* < 0.0001.

Previous studies showed that DMY administration prevents Aβ deposition and cognitive decline in APP/PS1 mice ^43^. However, none of these studies focused on the role of DMY administration in the interaction between cellular senescence and Aβ pathology. We next assessed whether elimination of senescent microglial cells through long-term intermittent senolytic treatment with DMY can affect SA-β-Gal activity and Aβ pathology in 5×FAD mice. Beginning at 3 months of age, female 5×FAD mice were treated with either placebo or DMY twice *per* week for 3 months. Spatial learning and memory were evaluated by testing the 5×FAD mice in the Y-maze at 6 months of age, whereas animals were euthanized at 6.5 months, with their brains processed for histological and biochemical analyses (Fig. 8e). Aβ plaque-associated SA-β-Gal activity in the cortex and hippocampus was diminished in the DMY-treated 5×FAD mice compared to the placebo-treated littermates (Fig. 8f, g and Supplementary Fig. 14c, d). Unlike the acute DMY treatment, there was a marked reduction in Aβ plaque load in the cortex and hippocampus of DMY-treated 5×FAD mice compared to placebo-treated (Fig. 8f, g and Supplementary Fig. 14c, d). Moreover, DMY administration reduced levels of Aβ_1-42_ and Aβ_1-40_ in serum, suggesting potential effects of the DMY senolytic treatment on Aβ production and/or clearance (Fig. 8h). In line with a reduction of IL6 level in response to DMY treatment (Fig. 8f, g and Supplementary Fig. 14c, d), the transcript level of the senescence biomarker p16^INK4A^ and various proinflammatory cytokines (TNF-α, IL6, IL1β) was significantly lower in the cortex and hippocampus of DMY-treated AD mice compared to those treated with placebo (Fig. 8i and Supplementary Fig. 14e). We also observed that SA-β-Gal positivity associated with activation of microglial and astrocytes were significantly reduced in the cortex and hippocampus of DMY-treated 5×FAD mice compared to placebo-treated (Supplementary Fig. 14f, g). Unsurprisingly, as compared with placebo-treated 5×FAD mice, those treated with DMY over the long-term performed significantly better in Y-maze test, grip strength assay and rotarod test (Fig. 8j-l). These data suggested that elimination of senescent microglial cells through DMY administration is sufficient to restore tissue homeostasis and physiological function, alleviating Aβ pathology and cognitive impairment in AD mice.

## Discussion

The identification of the first generation senolytics, ‘D + Q’, and their rapid transition from discovery to efficacy demonstration in animal models to clinical trials has inspired further and intensive senotherapeutic exploration. Most senotherapeutics reported to date are mechanistically described as being senolytics, selectively killing senescent cells by targeting key intracellular pathways associated with senescence-associated phenotypes particularly anti-apoptotic and pro-survival networks ^44, 45^. In contrast, senomorphics, which dampens the pathological SASP expression and extracellular secretion without eliminating senescent cells, may be as a similar or more effective and safer alternative to the reported senolytics ^6^. However, identification of mechanistic actions of senotherapeutics can be challenging, especially for agents that have multiple disease-improving roles. In this study, we revealed the potential of DMY as a senotherapeutic agent with dual functions, making it a promising agent for future intervention of age-related pathologies in geriatric medicine (Fig. 9).

**Fig. 9.**
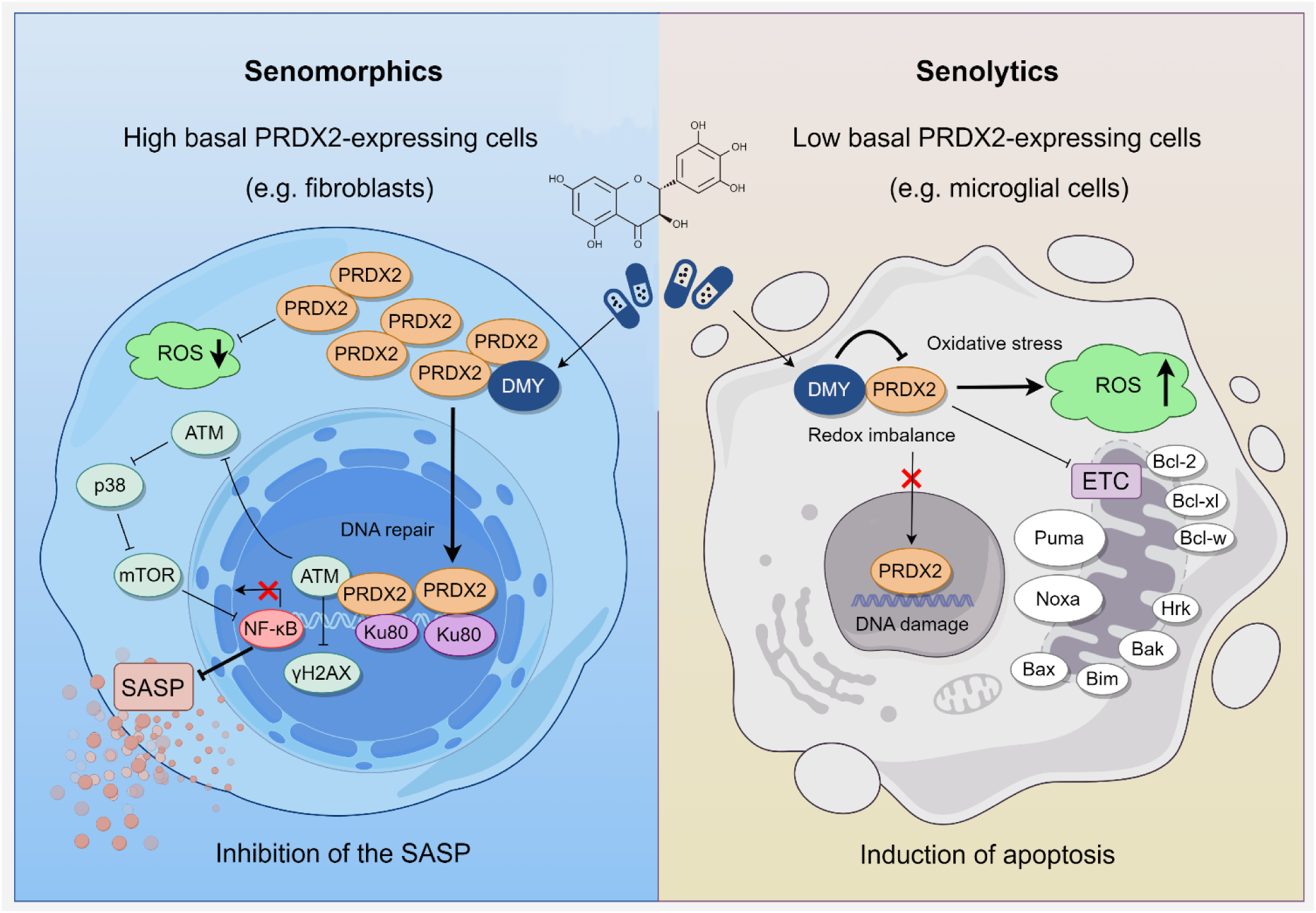
Schematic illustration of DMY as a senotherapeutic agent against senescent cells. For cell types of high-level basal expression of PRDX2, DMY targets this enzyme to facilitate DDR repair and inhibits the SASP, usually acting as a senomorphic agent. However, for cell types of low-level basal expression of PRDX2, DMY targets the oxidoreductase activity of PRDX2 and impairs mitochondrial function to cause senescent cell apoptosis. Thus, DMY can generate differential effects in a cell type-dependent manner. Through its senomorphic and/or senolytic action (s), PRDX2 holds a geroprotective potential to delay, prevent or ameliorate organismal aging and intervene in age-related pathologies.

Since senescent cells accumulate slowly in the course of lifespan, senolytics can be administered periodically, and supposed to prevent dysregulation of senescent cell-driven critical processes such as tissue repair and wound healing ^8^.

Senomorphics, which attenuate the deleterious and pathological SASP without inducing radical senolysis, may represent an optimal therapeutic option. Like senolytics, senomorphics have been demonstrated to extend both healthspan and lifespan in animals, although they may have side-effects. For instance, treatment with the senomorphic agent rapamycin, which remarkably alleviates age-associated dysfunctions in model organisms, largely resembling the senolytic drug navitoclax, frequently induces hyperlipidaemia, metabolic dysregulation, thrombocytopenia and impairs wound healing ^46^. In contrast, rutin, another natural senomorphic agent, represents an effective and potent therapeutic avenue to control age-related pathologies without inducing overall side-effects ^23^. Similarly, DMY administration to animals has been proved to be generally safe, without evident side-effects reported so far, another advantage of this agent.

The development of senotherapeutics to intervene age-related pathologies and achieve healthy longevity is progressing at an accelerated rate. The field of senotherapeutics will continue to expand with more effective agents and drug combinations identified, although many of them need to be tested in clinical trials. DMY is a polyphenol hydroxyl dihydroflavonol compound with high content in ampelopsis grossedentata, which is often used for herbal tea and traditional Chinese medicine ^47^. DMY generates anti-inflammatory, antibacterial and antiviral effects, without obvious cytotoxicity reported in normal cells, a fact indicating its relatively high bio-safety ^48^. Despite the poor water solubility and relative stability in low temperatures and weak acidic environments (pH 6.0), DMY can distribute in various tissues and penetrate the blood-brain barrier after oral administration in rats, a prominent advantage supporting to intervene various pathologies ^49^. However, the inherent mechanism that allows DMY to exert geroprotective effects remains unclear. We administered DMY at a concentration of 20 mg/kg to treat prematurely aged mice and observed markedly alleviated age-related symptoms, an effect presumably mediated by its senomorphic activity. In the 5xFAD mouse model that develops typical AD pathologies, DMY selectively depleted senescent microglial cells from Aβ plaques, mitigated Aβ-associated conditions and prevented cognitive decline in these AD mice, a process mainly correlated with the senolytic activity of DMY.

PRDX2 represents an antioxidant enzyme extensively distributed in various cell types including but not limited to neurons and vascular smooth muscle cells ^50^. Reported as one of the most effective scavenger proteins against ROS and hydrogen peroxide (H_2_O_2_), PRDX2 protect cells against oxidative stress and associated injuries ^50^. However, PRDX2 knockdown causes a decline of redox-stress response capacity (RRC) and accelerated senescence in fibroblasts, while *Prdx2*-mutant C. elegans also showed decreased RRC ^34^. Thus, PRDX2 is essential to maintain cellular homeostasis in response to redox-stress stimuli. In our study, the basal expression of PRDX2 appears to differ substantially between microglial cells and other cell types such as fibroblasts (and endothelial cells), a condition that allows cells to respond differentially to DMY treatment. When DMY was used against senescent fibroblasts, DMY interacts with PRDX2 and promotes its nuclear translocation to facilitate DDR and reduce ROS levels. However, when DMY was employed to treat senescent microglial cells, it interacts with PRDX2 and prevents it from nuclear translocation, causing elevated redox imbalance and resulting in cell apoptosis. Thus, the differential effects of PRDX2 or its function mode is largely dependent on cell types and basal intracellular level of PRDX2, while regardless of DMY concentrations (Fig. 9). Although it is not uncommon that a senotherapeutic agent can exert differential effects depending on certain factors, our study defines DMY as a typical dual function agent and highlights its special value when exploited to target the aging process of different organs in a geroprotective setting.

The 5xFAD transgenic mouse model is frequently employed in AD studies, as it recapitulates a number of AD-associated phenotypes with a relatively early onset and aggressive age-dependent progression ^51^. Besides developing Aβ-containing plaques alongside neuroinflammation by 2 months of age, as well as exhibiting neuronal decline by 4 months of age, which intensifies by 9 months of age, these mice manifest a broad spectrum of behavioral impairments, together defined as AD-related neuropathology ^52^. Physical motor problems, associated with agility and reflex movements, including deficiency in balance and coordination, and skeletal muscle function, also typically arise by 9 months of age ^53^. We administered DMY to these animals, and observed significantly improved motor skill, learning capacity and memory ability. These changes were accompanied by reduced senescent cell accumulation, Aβ deposition and expression of the SASP markers, as well as decreased positivity of microglial cells and astrocytes in hippocampus and cortex. Although short-term treatment with DMY reduced microglial cells, long-term intervention caused reduction of both microglial cells and astrocytes, a phenomenon that contributes to the overall strength of DMY in restraining chronic neuroinflammation developed in brain tissues of AD animals, as DMY holds anti-inflammation potential, or more specifically, anti-SASP capacity in an aged host. However, DMY appeared to be less effective against senescent oligodendrocytes or astrocytes in our study, suggesting a limitation that may compromise its therapeutic potential in AD-related settings.

Our study highlights that DMY, a natural flavonoid abundant in plants such as vine tea, grapes and bayberry, can generate senotherapeutic effects by acting as a senomorphic and/or senolytic agent. Data from preclinical studies suggests that pharmaceutical intervention with DMY is a regime to ameliorate age-related conditions through targeting senescence *per se*. As senescent cells contribute to a wide range of age-related pathologies, establishing the impact of DMY-mediated therapeutic effects in aged or disease animals provides an efficient modality to target senescence via differential mechanisms in a systemic manner and to eliminate the radical cause of multiple comorbidities, a major topic of future geriatric medicine.

## Methods

### Cell culture

The primary normal human prostate stromal cell line PSC27 and the breast fibroblast cell line HBF1203 were provided by Dr. Peter Nelson (Fred Hutchinson Cancer Center) and maintained in a complete stromal medium as described previously ^54^. The human umbilical vein endothelial cell line HUVEC was acquired from ATCC and cultured in DMEM medium, supplemented with 10% fetal bovine serum (FBS). The prostate cancer epithelial cell lines PC3, DU145 and LNCaP were from ATCC and routinely cultured in RPMI 1640 medium supplemented with 10% FBS. The prostate cancer epithelial line M12 was from Dr. Stephen Plymate (University of Washington). Furthermore, human brain tissue-derived cell lines HMC3, MO3.13 and SVG p12, and murine microglial cell line BV2 were from the National Collection of Authenticated Cell Cultures (China) and were regularly cultured in DMEM medium supplemented with 10% FBS. All cell lines underwent routine test for mycoplasma contamination and were authenticated using Short Tandem Repeat assays.

### Cell treatments

Cells were cultured until reaching approximately 80% confluence (CTRL) under the condition of 5% CO_2_ at 37°C, and subsequently treated with 50 μg/ml bleomycin for a duration of 6-12 hr. Following treatment, cells were briefly washed with phosphate buffered saline (PBS) and maintained for a period of 7-10 d to allow for cellular senescence, before subject to various analyses. The senotherapeutic agents screened in this study were commercially acquired from TargetMol L8200-Anti-aging Compound Library. To identify potential senotherapeutics, natural candidates (totally 50 in the NMA library) were tested each at 100 μM on the survival of 5.0 × 10^3^ non-senescent and senescent cells for 3 d (senolytics). Subsequently, appraisal of the effects of remaining agents (50 in the library, each applied at 50 μM) was performed to test the ability to inhibit the SASP (senomorphics). For those demonstrating significant potential as effective and safe senotherapeutics, further assessments were followed. The phytochemical agent DMY was tested across a concentration gradient 0 μM to 800 μM, with 100 μM identified as a fairly effective concentration for senomorphic activity in PSC27 cells, with 100 μM (starting from 50 μM) as an effective concentration for senolytic activity in HMC3 cells. Additionally, the small molecule inhibitor SB203580 targeting p38MAPK was utilized at a concentration of 20 μM rutin, while the PRDX2 inhibitor Conoidin A was applied at concentrations 1 μM to 4 μM to treat senescent cells in culture prior to cell lysis and expression assays.

### HPLC-QTOF-MS/MS analysis

The standard high-performance liquid chromatography (HPLC) coupled with quadrupole time-of-flight tandem mass spectrometry (HPLC-QTOF-MS/MS) was performed on a Nexera X2 LC-40 system (SHIMADZU), integrated with the AB SCIEX TripleTOF 6600 LC/MS/MS apparatus (SCIEX). The data were processed using Analyst TF software (version 1.7.1). Briefly, the chromatographic separation was conducted on an Accucore C30 column (2.6 μm, 250 × 2.1 mm, Thermo Fisher) at room temperature. The mobile phase consisted of 1% acetic acid in water (designated as solvent A) and 100% methanol (designated as solvent B). A multistep linear gradient was employed, beginning at 5% solvent B and progressing to 15% B within 5 min, reaching 35% B at 30 min, and 70% B at 45 min, sustaining 70% B until 49 min, followed by a transition to 100% B at 50 min. The system was held at 100% B for an additional 10 min before returning to 5% B by the 60-min mark. During both the initial and final gradient stages, the column underwent equilibration and maintenance or washing with 5% B for a duration of 10 min. The flow rate was maintained at 0.2 ml/min, with a sample injection volume of 10 μl. Detection was performed *via* the TripleTOF 6600 qTOF mass spectrometer (AB SCIEX), which utilized an electrospray ionization (ESI) interface in negative ion mode. The operating parameters of the ESI source included a capillary voltage of -4000 V, a drying gas flow rate of 60 (arbitrary units), nebulization gas pressure set at 60 psi, a capillary temperature of 650°C and a collision energy of 30 eV. Mass spectra were recorded over a mass-to-charge ratio (m/z) range of 50 to 1200, with an acquisition rate of 3 spectra per second. Data acquisition and subsequent analysis were conducted using Analyst TF software (AB SCIEX, version 1.7.1).

### Human proteome microarray

DMY coupled with biotin (DMY-Biotin) was synthesized and identified by NMR, which was performed by Wayen Biotechnologies (Shanghai, China). D-Biotin was used as a control. The human proteome microarray was also sourced and performed by Wayen, wherein the HuProt^TM^ 20K microarray (CDI Laboratories, USA) consists of 20,240 full-length human proteins. Microarrays were immersed in pre-cooled blocking buffer (5% BSA and 0.1% Tween 20 in PBST) for 5 min and treated with blocking buffer at room temperature for 1.5 hr with gentle agitation in dark. DMY-Biotin and D-Biotin were then diluted to a concentration of 10 μM in the blocking buffer and applied to the previously blocked proteome microarray, and incubated at room temperature for 1 hr. Microarrays were washed thrice with PBST, each for 5 min. Microarrays were then treated with a 0.1% Cy5-Streptavidin solution for 20 min at room temperature, followed by 3 additional washes with PBST, each lasting 5 min. The microarrays were subsequently scanned with a GenePix 4000B microarray scanner (Axon Instruments, USA) to visualize and record the results. Data analysis was performed utilizing GenePix^TM^ Pro v6.0 software (Axon Instruments, USA).

### DMY-Biotin-target protein pull-down and LC-MS/MS analysis

Pull-down experiments along with LC-MS/MS analyses were performed by Wayen Biotechnologies (Shanghai, China). Briefly, proteins were extracted from senescent HMC3 and PSC27 cells using immunoprecipitation-specific lysis buffer containing protease inhibitors (1:100). For pull-down, a Pierce^TM^ Spin Column was prepared by adding TBS, which was subsequently immobilized and labeled with 5 mM DMY-Biotin or D-Biotin (as a control) overnight at 4°C. To block the Pierce^TM^ Spin Column, a biotin blocking solution was added at room temperature. Following this, 1 mg of the extracted protein was incubated with the Pierce^TM^ Spin Column at 4°C for 2 hr, after which the column was washed 3 times with a wash buffer. After reduction in 10 mM DTT, alkylation in 40 mM IAA and protease digestion with a trypsin solution (Promega, 1:50), the pull-down proteins were extracted using 50 mM NH_4_HCO_3_, before peptide solution was transferred to Solid Phase Extraction Cartridge (MonoSpin C18, GL Sciences) for desalting and clean-up.

Samples obtained above were analyzed by a Orbitrap Fusion Lumos mass spectrometer equipped with a Nanospray Flex source (Thermo Fisher Scientific, USA). Sample was separated by a C_18_ column (75 μm × 20 cm) packed with Reprosil-Pur C18-AQ, 1.9 mm resin on a nanoflow HPLC Easy-nLC 1000 system (Thermo Fisher Scientific). Buffer A comprising 0.1% (v/v) formic acid in H_2_O and Buffer B comprising 0.1% (v/v) formic acid in 80% acetonitrile. The separation was achieved using gradient elution program at 300 nl/min at 50°C: 1%-5% B in 4 min; 5%-26% B in 42 min; 26%-40% B in 10 min; 40%–100% B in 4 min; 100% B in 5 min. MS scan was performed in positive ion mode with spray voltage at 1,900 V, with ion transfer tube temperature at 320°C. The acquisition of MS data was conducted by Data Dependent Acquisition (DDA) mode. The MS1 full scan (350-1800) was set at a resolution of 60,000 @ m/z 200, automatic gain control (AGC) target 4e5 and maximum IT 50 ms. Isolation window was set at 0.7 m/z. MS2 scans were generated by HCD fragmentation at a resolution of 15,000 @ m/z 200, AGC target 1e5 and maximum IT 22 ms. The normalized collision energy (NCE) was set at NCE 30%, and the dynamic exclusion time was 30 sec. The Sequest HT search algorithm in Proteome Discoverer 2.4 was used to search MS/MS spectra against a composite database comprised of all the UniProt human database (swissprot_human_20423_20240516.fasta). Minimum cutoff for peptide length was set at 6 amino acids, and maximum permissible missed cleavage was set at 2. Maximal FDR for peptide spectral match, proteins and site was set to 0.01.

### Co-IP and LC-MS/MS analysis

PSC27 cells were induced senescent by exposure to 50 μg/ml BLEO for 12 hr before treated with 100 μM DMY or vehicle. Following this, cells were harvested using immunoprecipitation (IP) lysis buffer (Beyotime, P0013J) supplemented with protease and phosphatase inhibitors (1:100), and then incubated overnight at 4°C with a RPDX2 antibody (1:100, Abclonal, A1919) or IgG antibody (negative control). The resulting cell lysates were incubated with Protein A/G Magnetic Beads at 4°C for 2 hr. The magnetic beads were subjected to 5 washes with the IP lysis buffer. Proteins captured by the beads were subsequently analyzed using silver staining or Western blot. Alternatively, immunoprecipitated proteins were separated from magnetic beads using 8 M urea and subsequently analyzed by LC-MS/MS. Peptides were resolved in 0.1% formic acid, with the BCA protein quantification kit employed to measure peptide concentration. Samples were separated utilizing an analytic column (75 μm × 15 cm, 3 μm, 100Å) (C^18^, Thermo Fisher Scientific) on a nanoflow HPLC Easy-nLC 1200 system (Thermo Fisher Scientific, USA), using a 120 min LC gradient at 300 nl/min. Buffer A comprising 0.1% (v/v) formic acid in H_2_O and Buffer B comprising 0.1% (v/v) formic acid in 80% acetonitrile. The solution gradient was as follows: 1%-5% B in 6 min; 5%-26% B in 88 min; 26%-40% B in 22 min; 40%–100% B in 5 min; 90% B in 5 min. MS analyses were conducted on a Q Exactive HF-X mass spectrometer (Thermo Fisher Scientific). MS scan was performed in positive ion mode with spray voltage at 1,900 V, with ion transfer tube temperature at 320°C. Xcalibur software was used to acquire profile spectrum data in data-dependent acquisition pattern (DDA). The MS1 full scan was set at a resolution of 60,000 @ m/z 200, AGC target 3e6 and maximum IT 50 ms by orbitrap mass analyzer (350-1800 m/z), followed by ‘top 15’ MS2 scans generated by HCD fragmentation at a resolution of 15,000 @ m/z 200, AGC target 1e5 and maximum IT 30 ms. Isolation window was set at 1.6 m/z. The normalized collision energy (NCE) was set at NCE 28%, with the dynamic exclusion time as 45 sec. Precursors with charge 1, 7, 8 and > 8 were excluded from MS2 analysis. All raw mass spectrometric data were analyzed using MaxQuant 2.0.3.0 against the human Swiss-Prot database comprising 20,423 sequences (2024). Carbamidomethyl of cysteine was selected as a fixed modification. Oxidized methionine, protein N-term acetylation, lysine acetylation, asparagine and glutamine (NQ) deamidation were selected as variable modifications. Enzyme specificity was set as trypsin. The tolerances of first search and main search (Andromeda search engine) for peptides were set at 20 ppm and 6 ppm, respectively. Minimum cutoff for peptide length was set at 7 amino acids, with maximum permissible missed cleavage set at 2. Maximal FDR for peptide spectral match, proteins and site was set to 0.01. A minimum of 2 sequence-unique peptides was required for identification. Feature matching between runs was done with a retention time window of 2 min, with the label-free quantitation (LFQ) function enabled. The MaxQuant peptide and protein quantification results from the “peptides.txt” and “proteinGroups.txt” files were imported into Perseus software (version 1.5.1.6) for further analysis. Statistical significance between the groups was evaluated using Student’s *t* tests (*P* < 0.05), Proteins were defined as differentially expressed if the ratios were ≥ 1.5 or ≤ 0.67 in the experiment group compared with control.

### Quantitative proteomics

Quantitative proteomics were performed by Wayen Biotechnologies (Shanghai, China). Approximately 200 μg total protein from cells was digested. After desalting and cleanup, peptides were analyzed by a Orbitrap Astral mass spectrometer equipped with a Nanospray Flex source (Thermo Fisher Scientific, USA). Samples were loaded and separated by a C^18^ column of ES906 (PepMap TM Neo UHPLC 150 μM X 15 cm, 2 μM) (Thermo Fisher Scientific). Mobile phase A: 0.1% (v/v) formic acid in water; Mobile phase B: 0.1% (v/v) formic acid in acetonitrile/water (80:20, v/v). The separation was achieved using gradient elution program: 96% mobile phase A at 2.5 µl/min for 0-4 min, 75% mobile phase A at 2.5 µl/min for 4-5.8 min, 65% mobile phase A at 2.5 µl/min for 5.8-6.2 min, 1% mobile phase A at 2.5 µl/min for 6.2-6.9 min. Separated peptides proceeded directly to MS for DIA analysis after ionizing by Easy-spray ion (ESI) source. The ion spray voltage was set to 2.0 kV, with the ion transfer tube temperature maintained at 290°C. Raw files obtained from Astral mass spectrometer were analyzed by DIA NN, and the database was Swissprot human protein database (20,423 entries, 2024). Quantitative analysis was performed at the MS2 level using peak areas. Peptide abundance was calculated by summing the peak areas of their respective MS2 fragment ions, with protein abundance calculated by summing the abundance of their respective peptides. Upon Student’s *t*-test analysis, differential proteins with *P* value < 0.05 and fold change > 1.2 were selected. Numerical proteomic datasets are organized as additional Supplementary Files 1-4.

### SPR assay

The binding affinity of DMY to recombinant human PRDX2, PRDX1 and PRDX6 was assessed at 22°C utilizing the Biacore T200 instrument equipped with CM5 sensor chips (GE Healthcare, BR100530). Briefly, recombinant human protein in acetate acid buffer (pH 4.5) was covalently immobilized on the sensor chips following activation by 40 mM EDC and 10 mM NHS in aqueous solution. Various concentrations of DMY, ranging from 1.563 μM to 50 μM, were prepared with running buffer (PBS, 5% DMSO and 0.5% Tween 20). The DMY compound at different concentrations was injected simultaneously, injection of DMY solution was performed for 60 sec, with a dissociation time of 60 sec in each binding cycle. Data processing and analysis were performed using Biacore 4000 and Biacore T200 evaluation software (GE healthcare).

### CETSA assay

PSC27 cells were collected, and frozen-thawed 3 times using liquid nitrogen. The lysates were diluted with PBS containing protease inhibitor cocktail and divided into 2 aliquots, with 1 aliquot being treated with DMY (200 μM) and the other aliquot as control (DMSO). Each lysate was incubated with DMY or DMSO for 30 min at room temperature, then equally aliquoted into several parts in EP tubes and heated individually at different temperatures (35°C to 65°C) for 15 min. The precipitated proteins were separated from the soluble fraction by centrifugation at 14,000 × g at 4°C for 10 min, and boiled at 95°C for 10 min. Samples were subsequently used for immunoblot analysis. For the different drug concentration gradient CETSA experiments, PSC27 cell lysates were collected and aliquoted into several parts in EP tubes, then incubated with different DMY concentrations (0-400 μM) for 30 min at room temperature. Samples were heated at 55°C for 15 min before soluble fractions were isolated and analyzed by Western blot.

### DARTS assay

PSC27 cells were collected and total protein was lysed on ice with RIPA buffer containing protease inhibitors. Lysates were then treated with different DMY concentrations (0-400 μM) at room temperature. After incubation for 1 hr at room temperature, 6 μg/ml pronase was added into the lysates for a further 10 min at 37°C. Reactions were stopped by 1×SDS-PAGE loading buffer and samples were analyzed by Western blot.

### PRDX2 activity measurement

The activity of recombinant human PRDX2 protein was determined as previously described ^38^. PRDX2 exhibits the ability to reduce hydrogen peroxide (H_2_O_2_). Briefly, recombinant human PRDX2 proteins were either utilized alone or mixed with different concentrations of DMY, and incubated in a solution of 50 mM HEPES-HCl (pH 7.0) containing 1 mM DTT at room temperature for 10 min. The enzymatic reaction commenced with the introduction of 5 μl H_2_O_2_ (final concentration 100 μM) and incubated for 5 min. The remaining concentration of H_2_O_2_ was quantified, serving as an indicator of inhibitory effects on PRDX2’s catalytic function. Conoidin A was used as a positive control.

### Animals and drug treatments

All animal experiments were performed in compliance with NIH Guide for the Care and Use of Laboratory Animals (National Academies Press, 2011) and the ARRIVE guidelines, and were approved by the Institutional Animal Care and Use Committee (IACUC) of Shanghai Institute of Nutrition and Health, Chinese Academy of Sciences.

### Ionizing radiation

Animals were maintained in a specific pathogen-free (SPF) facility, with a controlled ambient temperature ranging from 22 to 25°C and humidity set at 50%. The animals were subjected to a 12-hr light-dark cycle, with unrestricted access to food and ad libitum feeding. To induce *in vivo* senescence, C57BL/6J mice (Model Animal Research Center of Nanjing University) were treated twice at 3-4 months of age with a sub-lethal dose of whole-body irradiation (WBI) at 5 Gy. Following a one-week recovery period, the mice received intraperitoneal injections of 20 mg/kg DMY or a placebo, administered twice every week for 3 consecutive months. Functional assays were performed after the final injection, after which mice were euthanized. Tissues were collected for RNA extraction and frozen in OCT solution for subsequent cryo-sectioning, SA-β-Gal staining, immunohistochemistry and immunofluorescence analyses. All mice involved in the experiments were randomly assigned to either control or treatment groups, with both male and female animals recruited.

### Chemotherapy

NOD-obese diabetic and severe combined immunodeficiency (NOD-SCID) mice (males, Model Animal Research Center of Nanjing University) aged 6 to 8 weeks were housed in a specific pathogen-free (SPF) environment, in compliance with the animal care guidelines established by the SINH. Each experimental group consisted of 10 mice, with tumor xenografts being established subcutaneously on the hind flank following anesthesia induced by isoflurane inhalation. Human prostate stromal cells (PSC27) were mixed with cancer cells (PC3) in a ratio of 1:4, specifically 250,000 stromal cells mixed with 1,000,000 cancer cells, to create tissue recombinants prior to *in vivo* implantation, as previously described ^22^. Following a 2-week period, mice were randomly assigned to various treatment groups for preclinical interventions. Treatments included MIT (0.2 mg/kg) alone, DMY (20 mg/kg) alone, a combination of MIT and DMY, or a placebo control, all administered via subcutaneous injection on the first day of the 3rd, 5th and 7th weeks, culminating in a total of 3 cycles over the entire 8-week treatment regimen. Tumor progression was monitored weekly, with tumor volume (v) evaluated and calculated using the formula: v = (π/6) × ((l+w+h)/3)³, where length (l), width (w), and height (h) of tumors were measured. Animals were euthanized, with tumors freshly excised and either snap-frozen or fixed for preparation of formalin-fixed paraffin-embedded (FFPE) samples. These prepared sections were subsequently utilized for SA-β-Gal staining, immunohistochemistry and immunofluorescence staining.

### AD mouse model and treatment

The 5×FAD transgenic mice were provided by Dr. Jiawei Zhou (Center for Excellence in Brain Science and Intelligence Technology, Chinese Academy of Sciences). These mice were identified by PCR-based genotyping and used for pharmacological analysis at 3 months of age. Genotyping primers: APP forward 5’-AGGACTGACCACTCGACCAG-3’, reverse 5’-CGGGGGTCTAGTTCTGCAT-3’; PS1 forward 5’-AATAGAGAACGGCAGGAGCA-3’, reverse 5’-GCCATGAGGGCACTAATCAT-3’. A cohort of 14 3-month-old female 5×FAD transgenic mice was randomly assigned to receive either intraperitoneal injections of 20 mg/kg DMY or a placebo, administered twice weekly for 3 consecutive months. In a separate experiment, ten 12-month-old female 5×FAD transgenic mice were randomly assigned to undergo acute short-term treatment, receiving a single daily intraperitoneal injection of 20 mg/kg DMY or placebo for a consecutive 7 d period prior to tissue collection. Upon completion of functional assays, mice were anesthetized using isoflurane and subsequently subjected to transcardial perfusion with PBS followed by 4% paraformaldehyde in PBS. The fixed and sucrose-cryoprotected brain tissues were sectioned into 10 μm sections for various staining.

### Physical function measurements

All measurements were performed at least 5 days after the last dose of DMY treatment.

### Y-maze test

The Y-maze spontaneous alternation was employed to evaluate short-term spatial memory using an opaque Perspex Y-maze (31 cm long, 5 cm wide and 10 cm high walls). Each arm had markers of different colors as distinct visual cues. Each animal was placed in turn in arm A of the Y-maze and allowed to explore freely for 5 min, with arm entries, alternations and walking distance recorded visually to calculate the percentage of the alternation behavior with the following formula: Alternation % = [Number of Alternations/ (Total number of arm entries - 2)] x 100. Spontaneous alternation (%) was defined as a successive entry into 3 different arms, on overlapping triplet sets.

### Grip strength test

The forelimb grip strength (N-F/kg) was evaluated by employing a grip strength meter (Shanghai XinRuan). To ensure the accuracy and reliability of results, the test was conducted over 10 trials. In each trial, mice were carefully positioned. Upon completion of all trials, results were averaged to obtain a comprehensive assessment of the forelimb grip strength.

### Rotarod test

The Rotarod test was utilized to assess motor coordination and balance with an advanced accelerating RotaRod system (Shanghai XinRua). Mice were trained on the RotaRod for a consecutive 2 d at a speed of 4 rpm for a duration of 200 sec. Subsequently, testing was conducted on the 3rd, 4th, 5th and 6th days. On the test day, mice were placed onto the RotaRod at an initial speed of 4 rpm/min. The rotating speed was gradually increased from 4 to 40 rpm/min over a 5 min interval. To accurately monitor the performance of the mice, a timer was employed to precisely record the moment when each mouse either dropped off the RotaRod or clung to the rod and completed a full passive rotation.

### Statistical analysis

Statistical analyses were predominantly conducted using GraphPad Prism version 8.0. A two-tailed unpaired Student’s *t*-test with Welch’s correction was used to assess statistically significant differences between 2 groups. One-way ANOVA with Tukey’s post-hoc comparison was used for multiple comparisons. Investigators were blinded to the allocation during the majority of experiments and outcome assessments. Animals were assigned to experimental groups based on their baseline body weight to ensure comparable body weights across groups, with randomization occurring only within groups. The determination of sample size was derived from prior experiments, so that statistical power analysis was not performed. Values are presented as mean ± SD unless otherwise indicated, with *P* < 0.05 considered significant. All data in this study were derived from independent biological replicates.

### Reporting summary

Further information on research design is available in the Nature Portfolio Reporting Summary linked to this article.

## Data availability

Data supporting the plots within this paper and other findings of this study are available from the corresponding author upon reasonable request. The RNA-seq data generated in the present study have been deposited in the NCBI Gene Expression Omnibus (GEO) database under accession code GSE190280 (human stromal cells), GSE275834 (human microglial cells) and GSE291050 (human stromal cells in mouse xenografts during preclinical treatment). The mass spectrometry proteomics data have been deposited to the ProteomeXchange Consortium *via* the PRIDE partner repository iProX (project ID no. IPX0011774000) with the ProteomeXchange dataset identifier PXD063165. Source Data are provided with this paper.

## Acknowledgements

We are grateful to the members of Sun laboratory for reagents, comments and other contributions to this project. The work was supported by grants from the National Natural Science Foundation of China (NSFC) (82471622, 82201737 to Q.X.); China Postdoctoral Science Foundation (BX20220323, 2021M703357 to Q.X.); the Strategic Priority Research Program of Chinese Academy of Sciences (XDB39010500 to Y.S.); National Key Research and Development Program of China (2020YFC2002800 to Y.S.), National Natural Science Foundation of China (NSFC) (82130045, 82350710221 and 82571777 to Y.S.); Shanghai Municipal Science and Technology Commission Excellent Academic Leader Program (20XD1404300 to Y.S.) and Taishan Scholar Project of Shandong Province (tstp20230634 Y.S.).

## Author contributions

Q.X. and Y.S. conceived this study, designed the experiments and orchestrated the project. Q.X. and G.L. performed most of the *in vitro* assays and *in vivo* experiments.

Q.X. analyzed data and wrote part of the manuscript. H.Z. and Z.J. helped with cell culture, drug treatment, sample preparation and proteomic analysis of human senescent cells. X.G. conducted SPR experiments on DMY interactions with PRDX1, PRDX2 and PRDX6. Z.L. assisted with MS analysis of the small molecule DMY.

L.G.P.L.P and J.L.K. delivered constructive advice. G.Z. provided technical support and constructive feedback. Y.S. supervised the study, provided funding support and conceptual input, and finalized the manuscript. All authors critically read and commented on the final manuscript.

## References

1. Sun, Y., Li, Q. & Kirkland, J.L. Targeting senescent cells for a healthier longevity: the roadmap for an era of global aging. Life Med 1, 103–119 (2022).

2. Lopez-Otin, C., Blasco, M.A., Partridge, L., Serrano, M. & Kroemer, G. Hallmarks of aging: An expanding universe. Cell 186, 243–278 (2023).

3. Gorgoulis, V. et al. Cellular Senescence: Defining a Path Forward. Cell 179, 813–827 (2019).

4. Song, S., Tchkonia, T., Jiang, J., Kirkland, J.L. & Sun, Y. Targeting Senescent Cells for a Healthier Aging: Challenges and Opportunities. Adv Sci (Weinh) 7, 2002611 (2020).

5. Gasek, N.S., Kuchel, G.A., Kirkland, J.L. & Xu, M. Strategies for Targeting Senescent Cells in Human Disease. Nat Aging 1, 870–879 (2021).

6. McHugh, D., Duran, I. & Gil, J. Senescence as a therapeutic target in cancer and age-related diseases. Nat Rev Drug Discov 24, 57–71 (2025).

7. Sun, Y., Li, Q. & Kirkland, J.K. Targeting Senescent Cells for a Healthier Longevity: the Roadmap for an Era of Global Aging. Life Medicine 1 (2022).

8. Zhang, L., Pitcher, L.E., Prahalad, V., Niedernhofer, L.J. & Robbins, P.D. Targeting cellular senescence with senotherapeutics: senolytics and senomorphics. Febs J (2022).

9. Lim, H., Park, H. & Kim, H.P. Effects of flavonoids on senescence-associated secretory phenotype formation from bleomycin-induced senescence in BJ fibroblasts. Biochem Pharmacol 96, 337–348 (2015).

10. Knopman, D.S. et al. Alzheimer disease. *Nat Rev Dis Primers* **7**, 33 (2021).

11. Gonzales, M.M. et al. Senolytic therapy in mild Alzheimer’s disease: a phase 1 feasibility trial. Nat Med 29, 2481–2488 (2023).

12. Cummings, J., Ritter, A. & Zhong, K. Clinical Trials for Disease-Modifying Therapies in Alzheimer’s Disease: A Primer, Lessons Learned, and a Blueprint for the Future. J Alzheimers Dis 64, S3–S22 (2018).

13. Ng, P.Y., Zhang, C., Li, H. & Baker, D.J. Senescence Targeting Methods Impact Alzheimer’s Disease Features in 3xTg Mice. J Alzheimers Dis (2024).

14. Bussian, T.J. et al. Clearance of senescent glial cells prevents tau-dependent pathology and cognitive decline. Nature 562, 578–582 (2018).

15. Musi, N. et al. Tau protein aggregation is associated with cellular senescence in the brain. Aging cell 17, e12840 (2018).

16. Qian, J. et al. Dihydromyricetin attenuates D-galactose-induced brain aging of mice via inhibiting oxidative stress and neuroinflammation. Neuroscience letters 756, 135963 (2021).

17. Zhang, J. et al. Recent Update on the Pharmacological Effects and Mechanisms of Dihydromyricetin. Front Pharmacol 9, 1204 (2018).

18. Wang, Y. et al. Recent update on application of dihydromyricetin in metabolic related diseases. Biomed Pharmacother 148, 112771 (2022).

19. Nie, H. et al. Molecular mechanisms and promising role of dihydromyricetin in cardiovascular diseases. Physiol Res 71, 749–762 (2022).

20. Zhu, Y. et al. Past and Future Directions for Research on Cellular Senescence. Cold Spring Harb Perspect Med 14 (2024).

21. Sun, Y. et al. Treatment-induced damage to the tumor microenvironment promotes prostate cancer therapy resistance through WNT16B. Nat Med 18, 1359–1368 (2012).

22. Dou, X. et al. PDK4-dependent hypercatabolism and lactate production of senescent cells promotes cancer malignancy. Nat Metab 5, 1887–1910 (2023).

23. Liu, H. et al. Rutin is a potent senomorphic agent to target senescent cells and can improve chemotherapeutic efficacy. Aging cell 23, e13921 (2024).

24. Zhang, B.Y. et al. The senescence-associated secretory phenotype is potentiated by feedforward regulatory mechanisms involving Zscan4 and TAK1. Nat Commun 9, 1723 (2018).

25. Wei, C. et al. Dihydromyricetin Enhances Intestinal Antioxidant Capacity of Growing-Finishing Pigs by Activating ERK/Nrf2/HO-1 Signaling Pathway. Antioxidants (Basel) 11 (2022).

26. Tse, A., et al. Improving the solubility of pseudo-hydrophobic Alzheimer’s Disease medicinal chemicals through co-crystal formulation. *bioRxiv* (2023).

27. Hou, L. et al. Dihydromyricetin resists inflammation-induced muscle atrophy via ryanodine receptor-CaMKK-AMPK signal pathway. Journal of cellular and molecular medicine 25, 9953–9971 (2021).

28. Fan, X. et al. Dihydromyricetin promotes longevity and activates the transcription factors FOXO and AOP in Drosophila. Aging 13, 460–476 (2020).

29. Keshava Prasad, T.S., et al. Human Protein Reference Database--2009 update. Nucleic Acids Res 37, D767–772 (2009).

30. Maglott, D., Ostell, J., Pruitt, K.D. & Tatusova, T. Entrez Gene: gene-centered information at NCBI. Nucleic Acids Res 39, D52–57 (2011).

31. UniProt, C. The Universal Protein Resource (UniProt) in 2010. Nucleic Acids Res 38, D142–148 (2010).

32. Sun, Z. et al. Dihydromyricetin alleviates doxorubicin-induced cardiotoxicity by inhibiting NLRP3 inflammasome through activation of SIRT1. Biochem Pharmacol 175, 113888 (2020).

33. Zhang, M. & Tang, Z. Therapeutic potential of natural molecules against Alzheimer’s disease via SIRT1 modulation. Biomed Pharmacother 161, 114474 (2023).

34. Meng, J. et al. Redox-stress response resistance (RRR) mediated by hyperoxidation of peroxiredoxin 2 in senescent cells. Sci China Life Sci 66, 2280–2294 (2023).

35. Singh, J.K. et al. Zinc finger protein ZNF384 is an adaptor of Ku to DNA during classical non-homologous end-joining. Nat Commun 12, 6560 (2021).

36. Lu, D. et al. Nuclear GIT2 is an ATM substrate and promotes DNA repair. Molecular and cellular biology 35, 1081–1096 (2015).

37. Wang, T. et al. Pan-cancer analysis of the oncogenic effects of G-protein-coupled receptor kinase-interacting protein-1 and validation on liver hepatocellular carcinoma. Adv Clin Exp Med 32, 1139–1147 (2023).

38. Chen, X. et al. Celastrol induces ROS-mediated apoptosis via directly targeting peroxiredoxin-2 in gastric cancer cells. Theranostics 10, 10290–10308 (2020).

39. Zhang, Y. et al. S-nitrosylation of the Peroxiredoxin-2 promotes S-nitrosoglutathione-mediated lung cancer cells apoptosis via AMPK-SIRT1 pathway. Cell Death Dis 10, 329 (2019).

40. Cacace, J. et al. Poor glycaemic control in type 2 diabetes compromises leukocyte oxygen consumption rate, OXPHOS complex content and neutrophil-endothelial interactions. Redox Biol 81, 103516 (2025).

41. Chen, F. et al. Targeting SPINK1 in the damaged tumour microenvironment alleviates therapeutic resistance. Nat Commun 9, 4315 (2018).

42. Xu, Q. et al. Targeting amphiregulin (AREG) derived from senescent stromal cells diminishes cancer resistance and averts programmed cell death 1 ligand (PD-L1)-mediated immunosuppression. Aging cell 18, e13027 (2019).

43. Feng, J. et al. Dihydromyricetin inhibits microglial activation and neuroinflammation by suppressing NLRP3 inflammasome activation in APP/PS1 transgenic mice. CNS Neurosci Ther 24, 1207–1218 (2018).

44. Yang, Y. et al. A BCL-xL/BCL-2 PROTAC effectively clears senescent cells in the liver and reduces MASH-driven hepatocellular carcinoma in mice. Nat Aging 5, 386–400 (2025).

45. Du, K. et al. Targeting senescent hepatocytes for treatment of metabolic dysfunction-associated steatotic liver disease and multi-organ dysfunction. Nat Commun 16, 3038 (2025).

46. Song, S., Lam, E.W., Tchkonia, T., Kirkland, J.L. & Sun, Y. Senescent Cells: Emerging Targets for Human Aging and Age-Related Diseases. Trends in biochemical sciences 45, 578–592 (2020).

47. Zhang, R. et al. Strategic developments in the drug delivery of natural product dihydromyricetin: applications, prospects, and challenges. Drug Deliv 29, 3052–3070 (2022).

48. Sun, C.C., Li, Y., Yin, Z.P. & Zhang, Q.F. Physicochemical properties of dihydromyricetin and the effects of ascorbic acid on its stability and bioavailability. J Sci Food Agric 101, 3862–3869 (2021).

49. Fan, K.J., Yang, B., Liu, Y., Tian, X.D. & Wang, B. Inhibition of human lung cancer proliferation through targeting stromal fibroblasts by dihydromyricetin. Mol Med Rep 16, 9758–9762 (2017).

50. Balasubramanian, P. et al. Implications and progression of peroxiredoxin 2 (PRDX2) in various human diseases. Pathol Res Pract 254, 155080 (2024).

51. Weible, A.P., Stebritz, A.J. & Wehr, M. 5XFAD mice show early-onset gap encoding deficits in the auditory cortex. Neurobiol Aging 94, 101–110 (2020).

52. Lin, C.J. et al. Mast cell deficiency improves cognition and enhances disease-associated microglia in 5XFAD mice. Cell Rep 42, 113141 (2023).

53. Padua, M.S., Guil-Guerrero, J.L. & Lopes, P.A. Behaviour Hallmarks in Alzheimer’s Disease 5xFAD Mouse Model. Int J Mol Sci 25 (2024).

54. Sun, Y. et al. Treatment-induced damage to the tumor microenvironment promotes prostate cancer therapy resistance through WNT16B. Nat Med 18, 1359–1368 (2012).

