## Supplementary Figures and Tables for "The natural flavonoid dihydromyricetin targets senescent cells *via* PRDX2 and alleviates age-related diseases"

### Supplementary Fig. 1

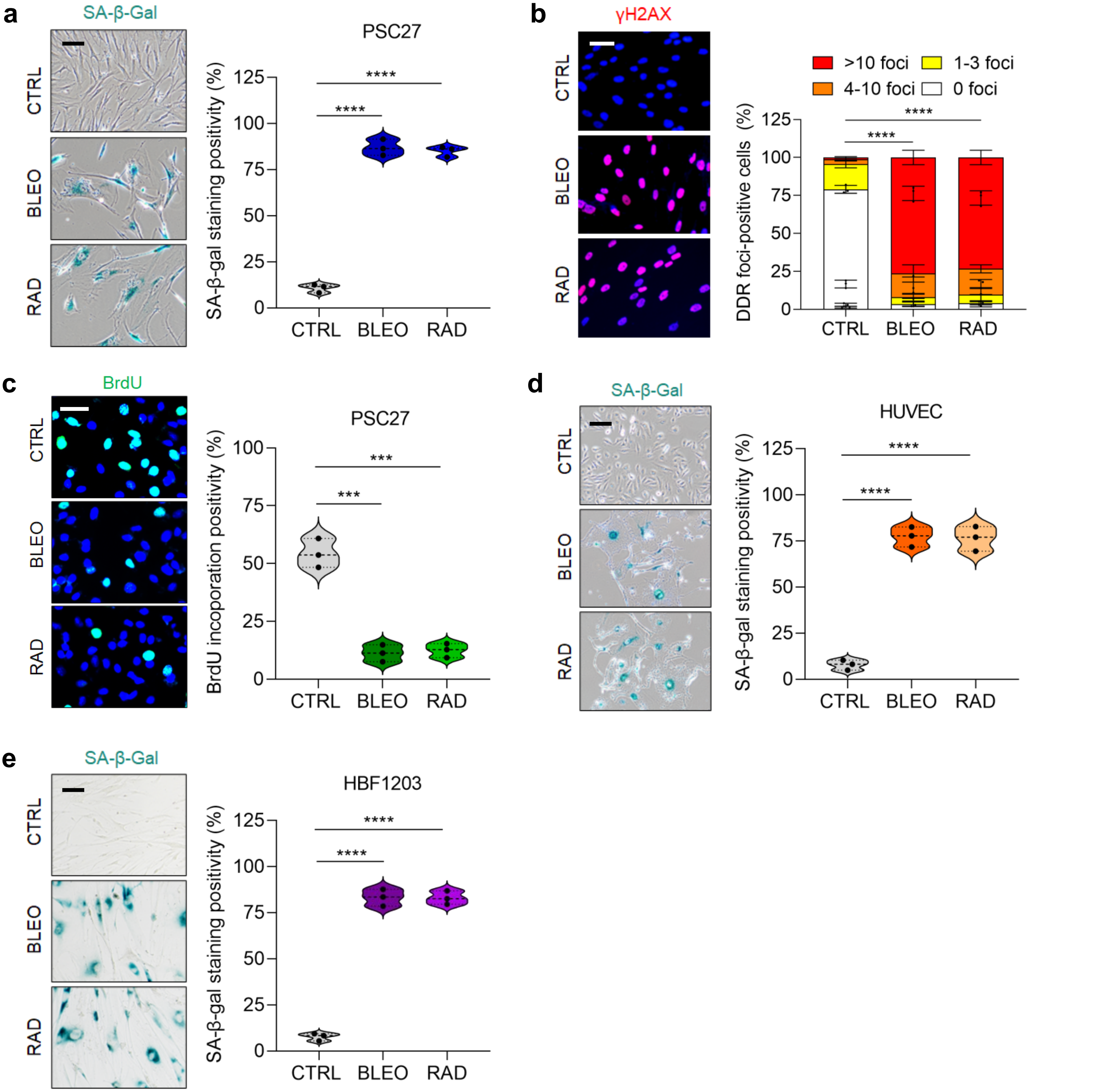

Supplementary Fig. 2

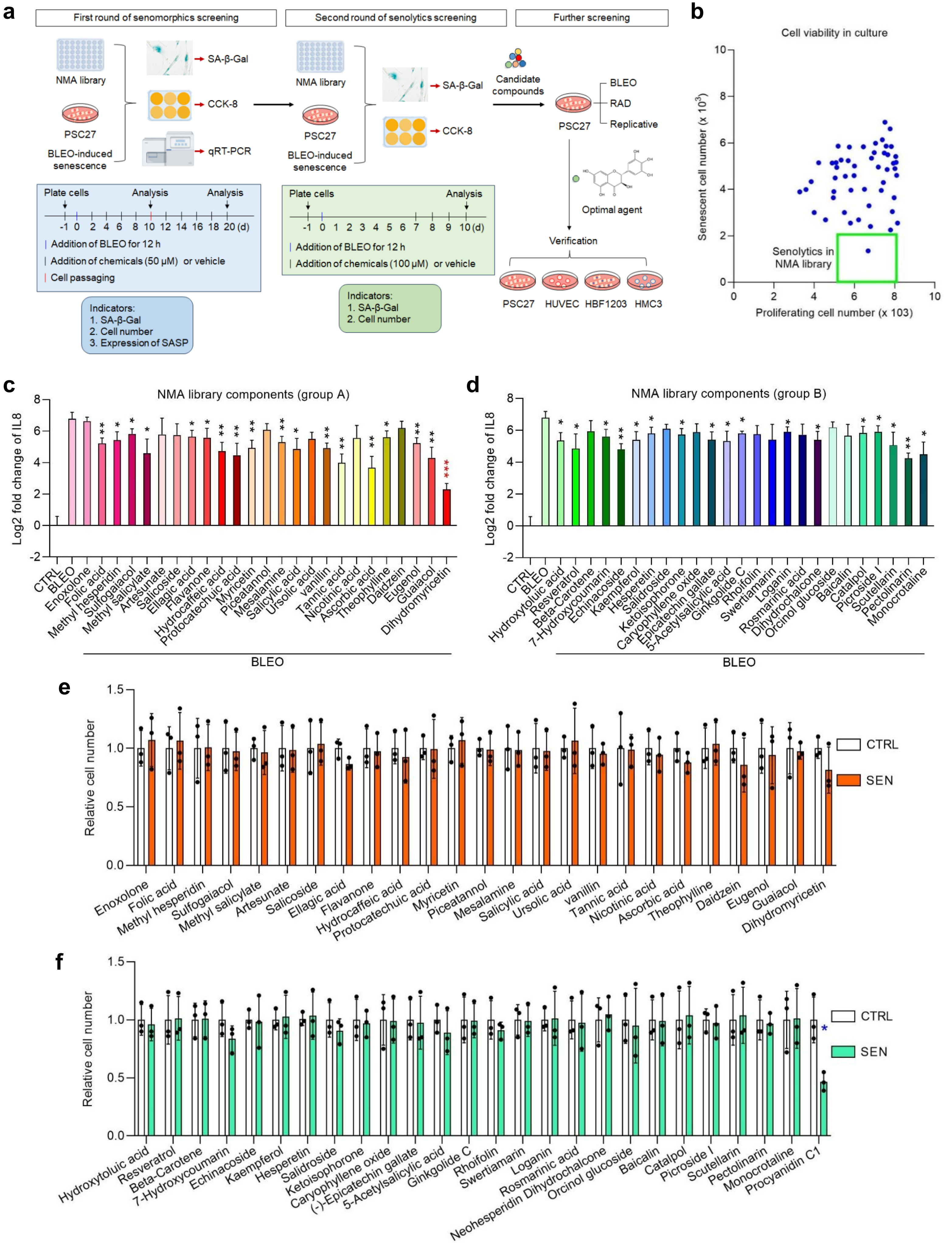

**Supplementary Fig. 3**

**a**

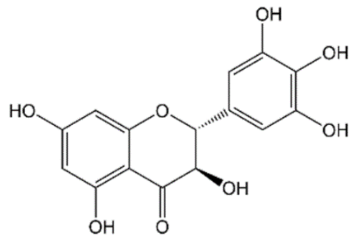


Dihydromyrcetin (CAS:27200-12-0)

**b**

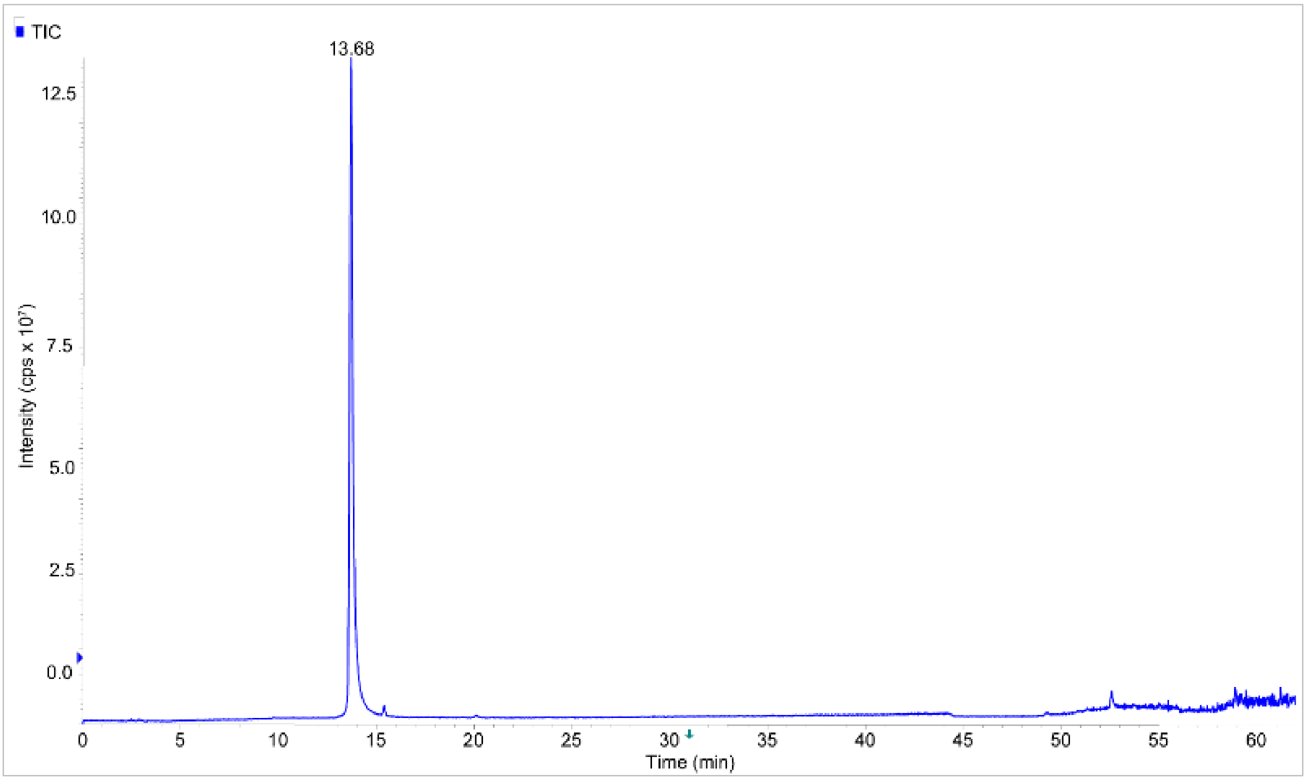


**c**

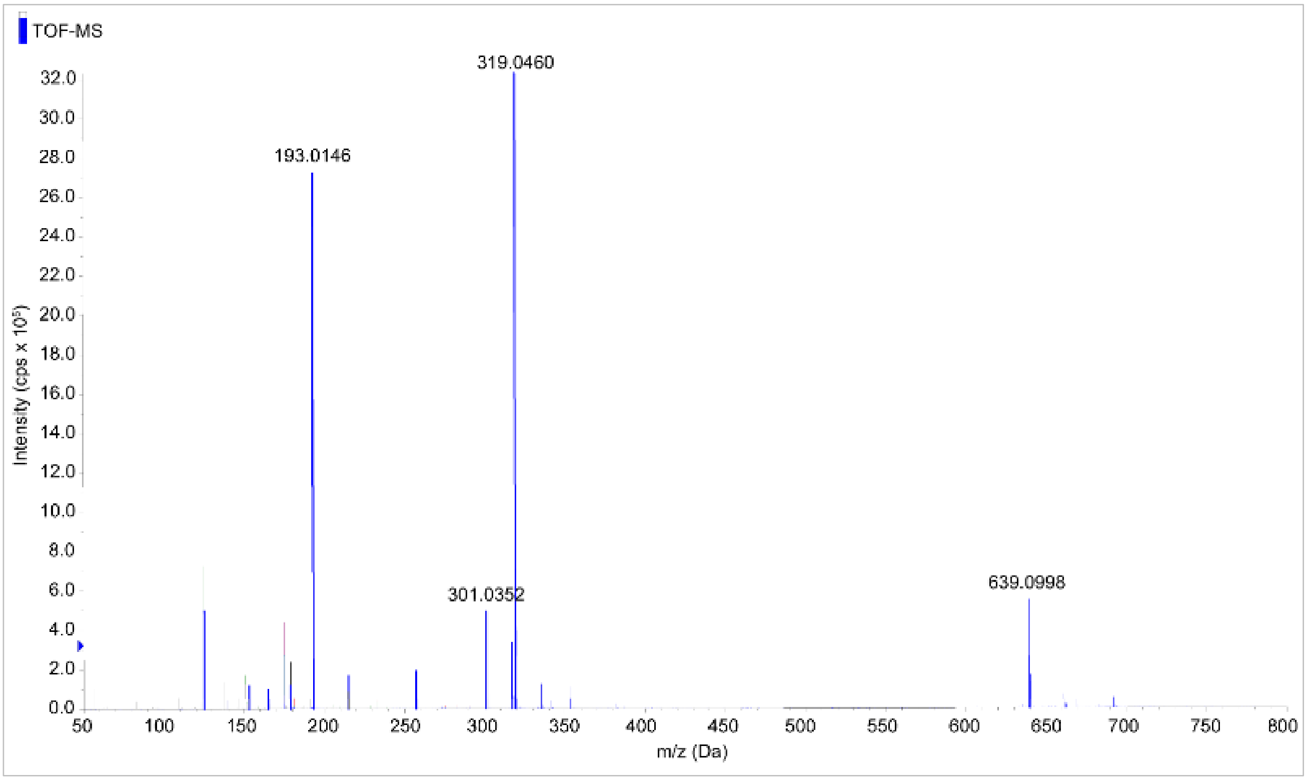

### Supplementary Fig. 4

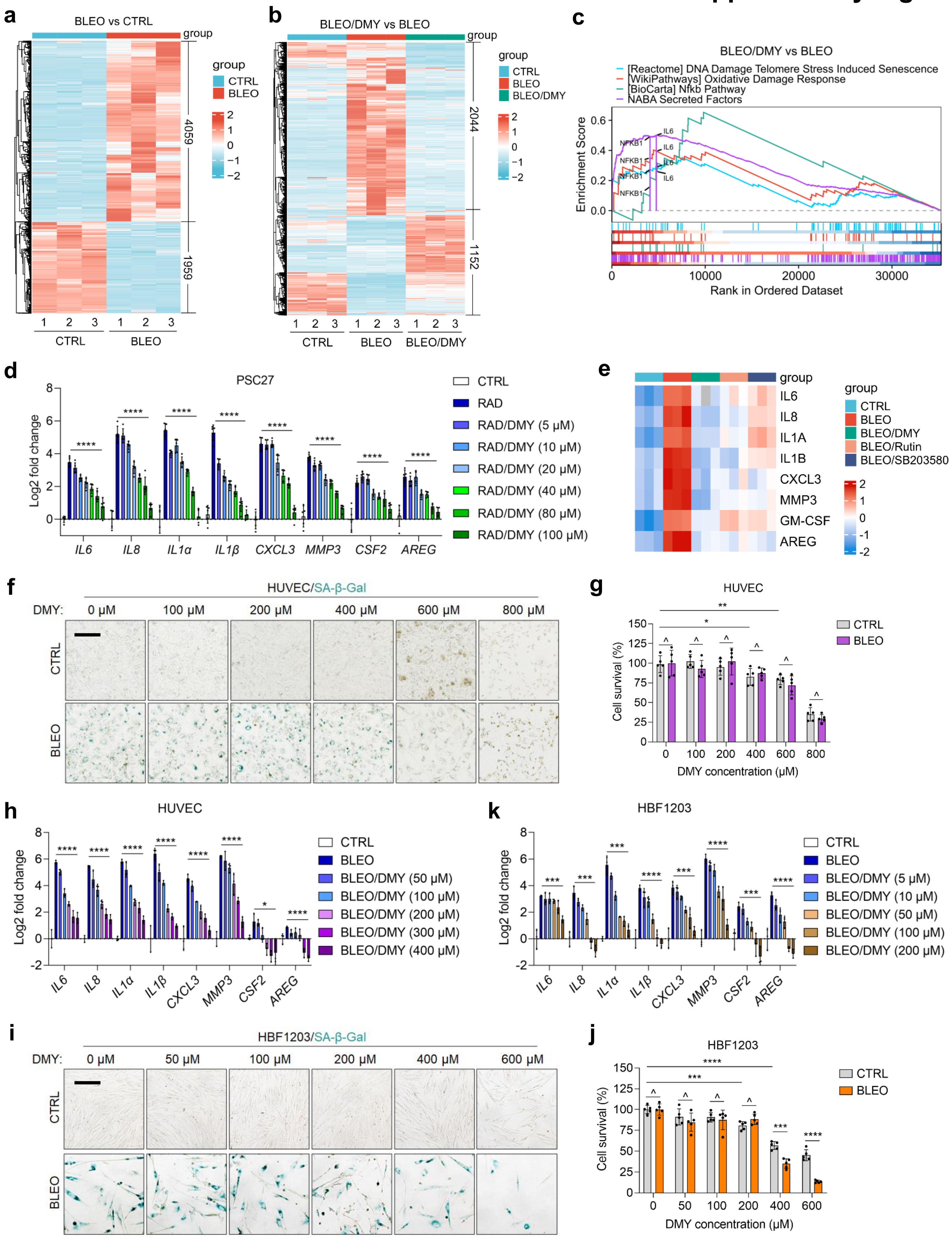

### Supplementary Fig. 5

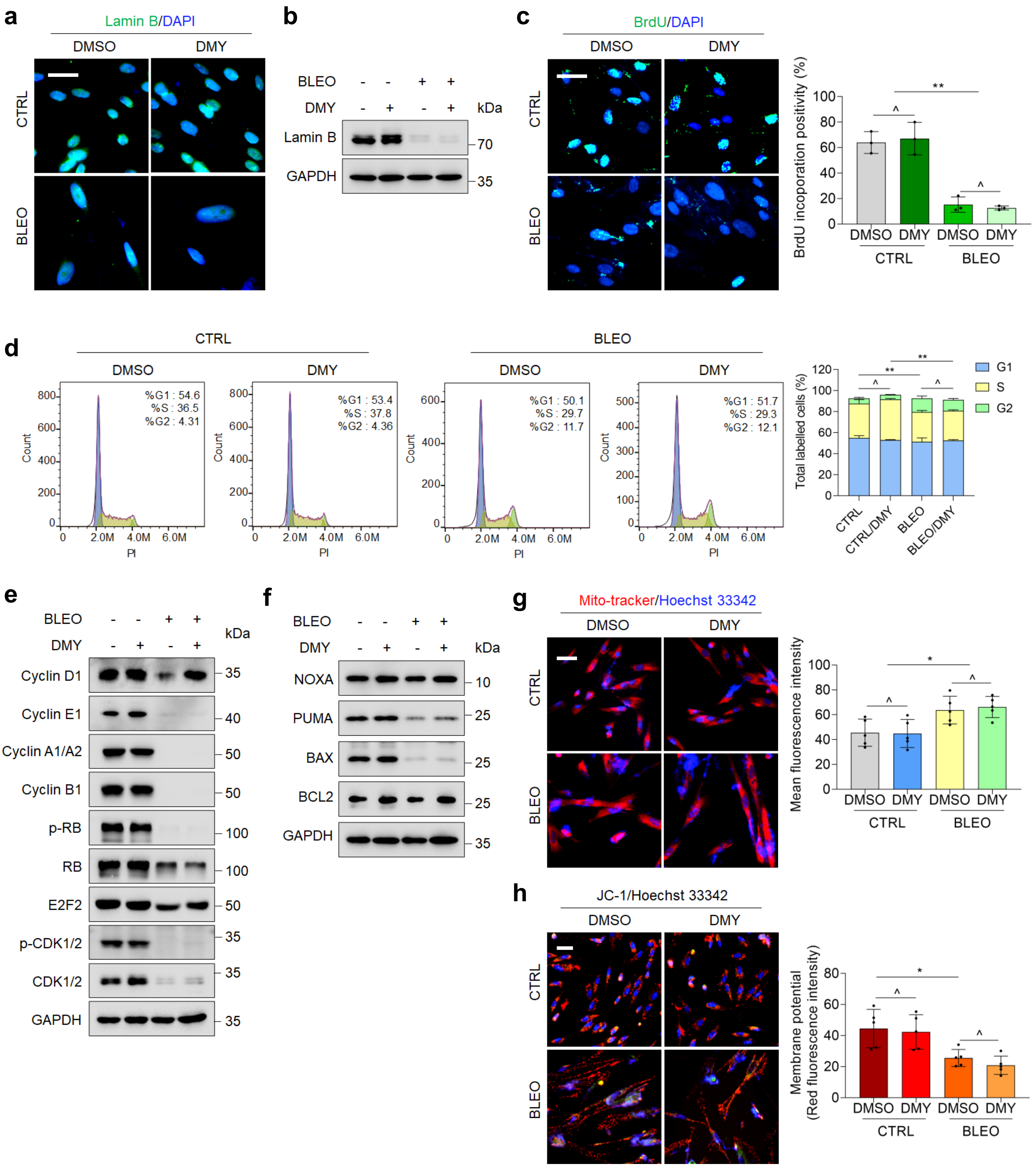

### Supplementary Fig. 6

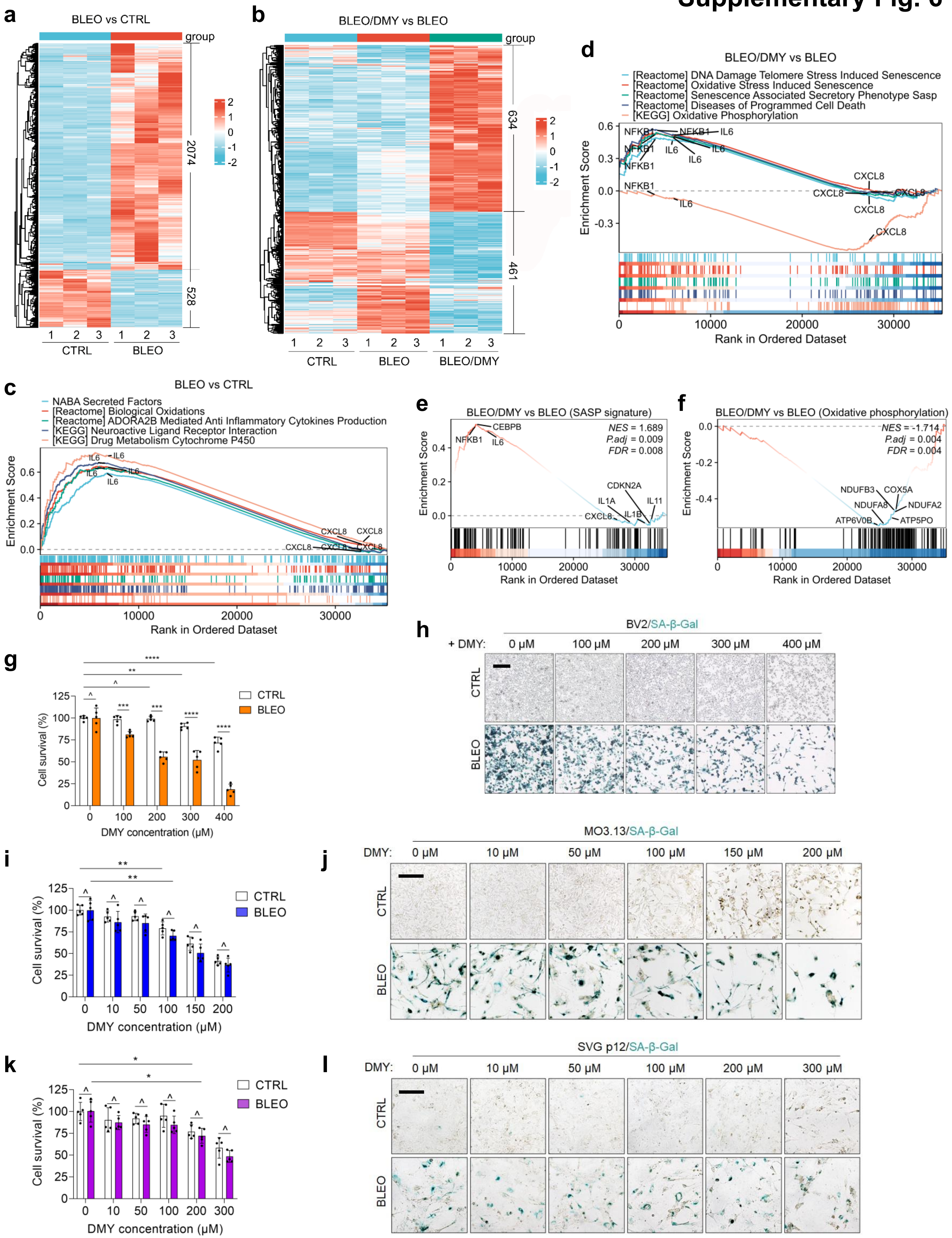

### Supplementary Fig. 7

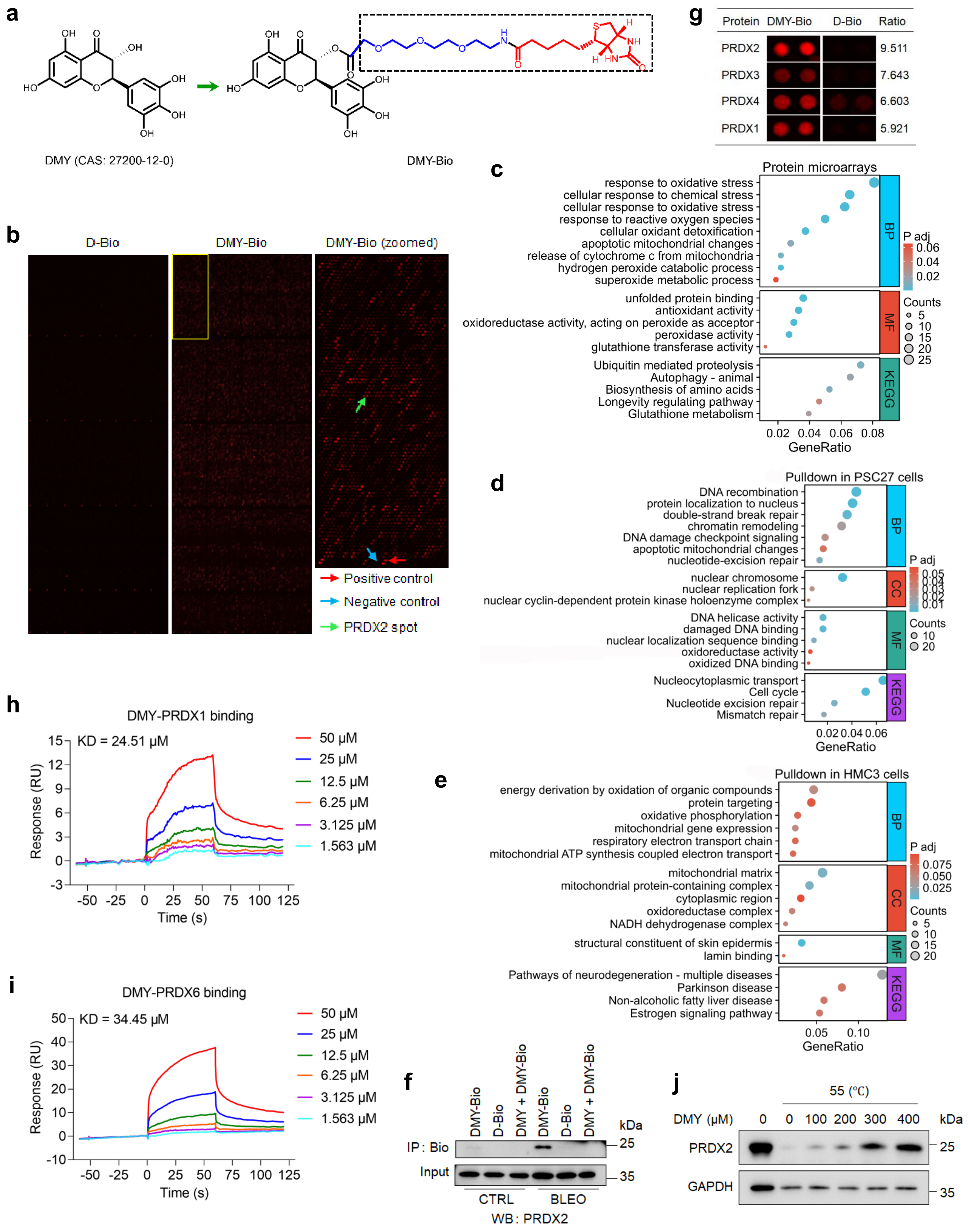

### Supplementary Fig. 8

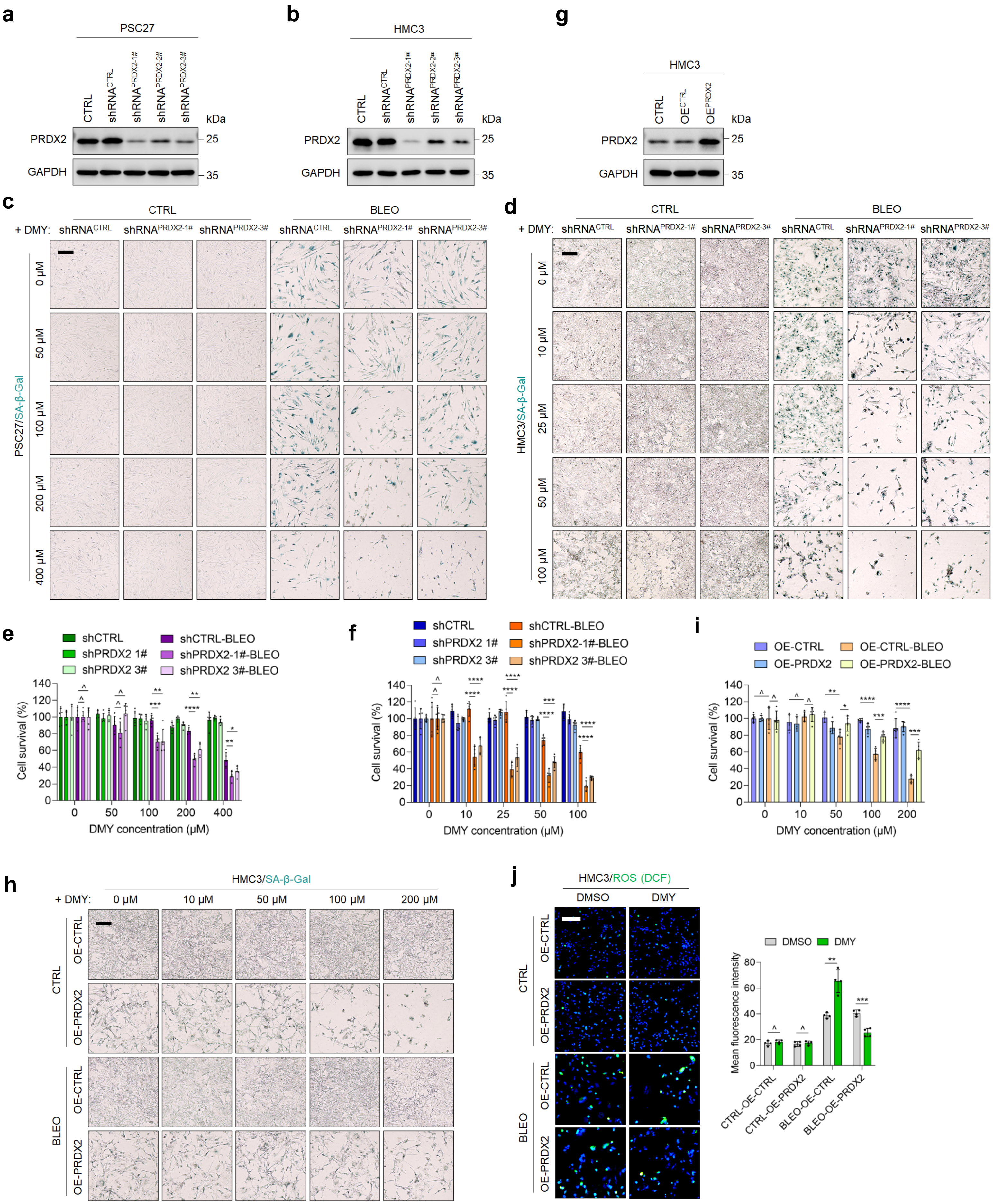

### Supplementary Fig. 9

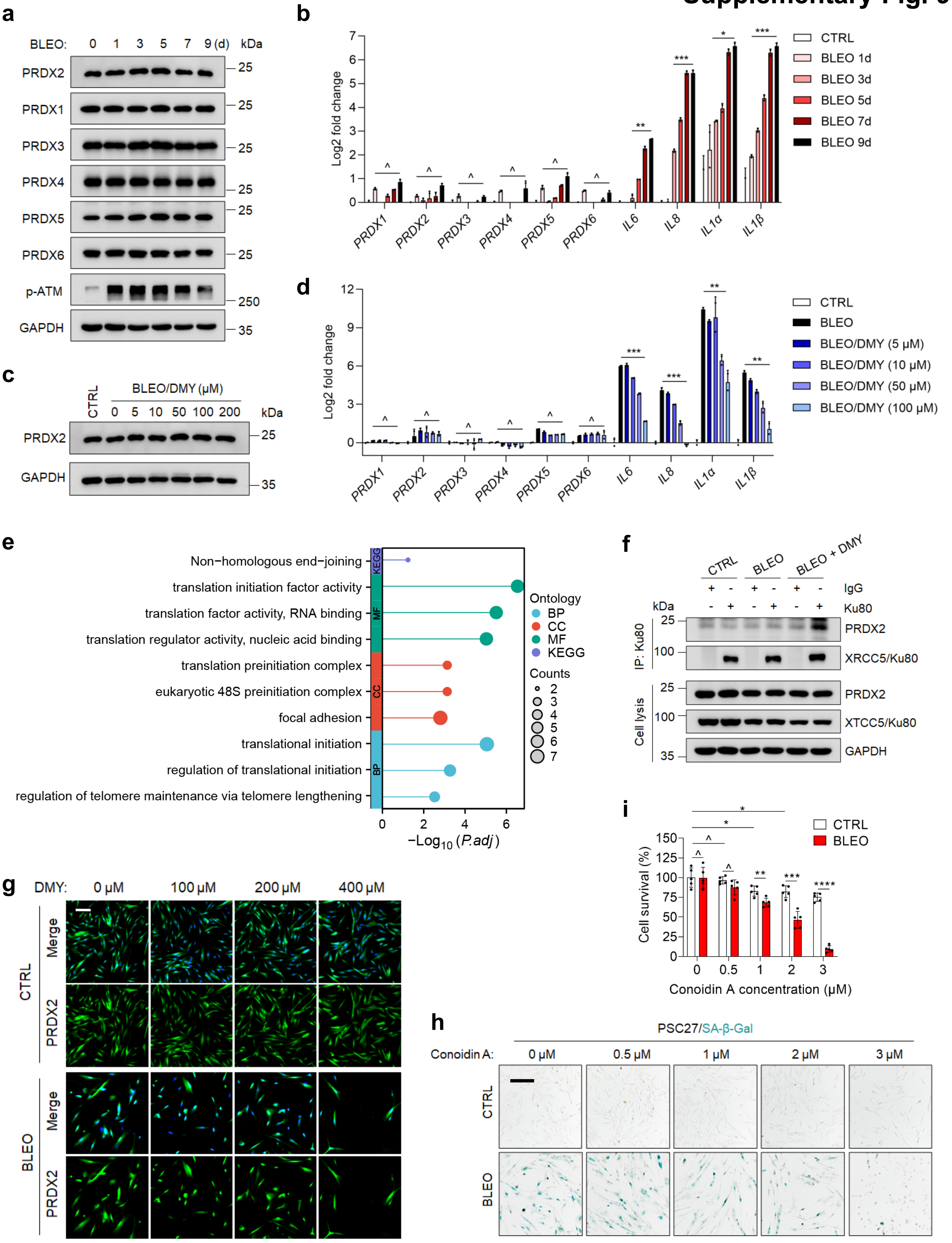

### Supplementary Fig. 10

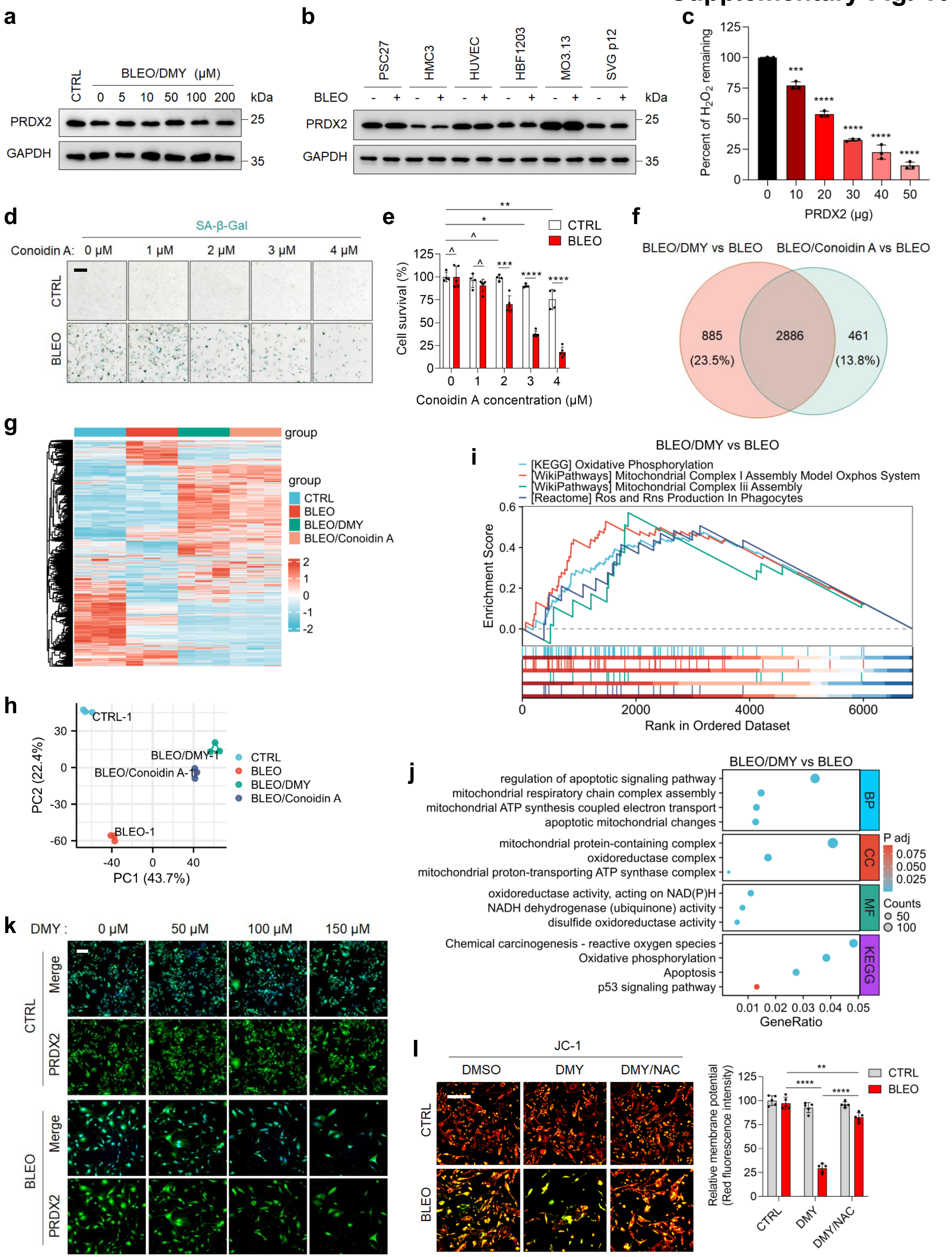

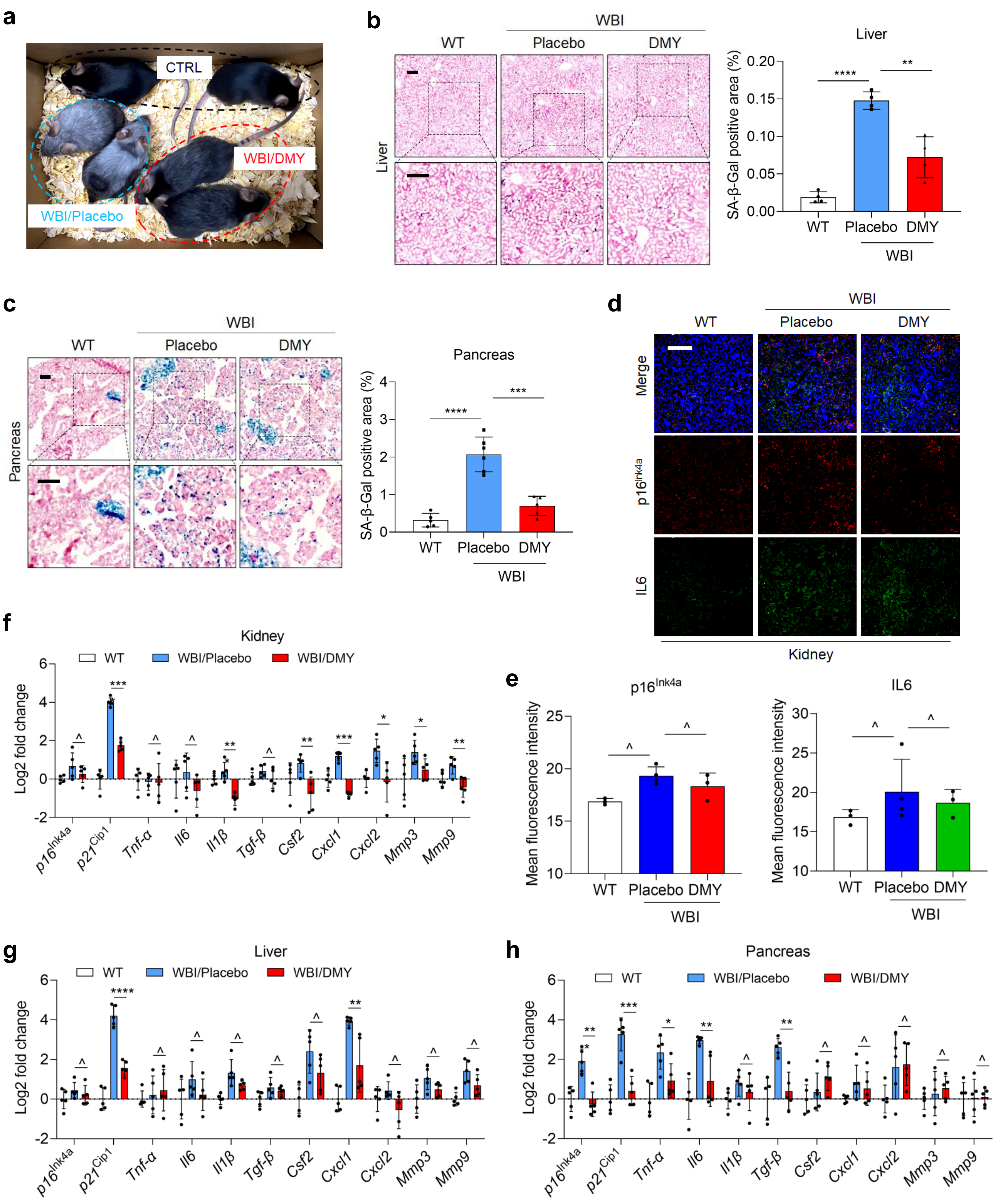
a
CTRL
WBI/DMY
WBI/Placebo
b
WBI
WT
Placebo
DMY
Liver
Liver
SA-β-Gal positive area (%)
c
WBI
WT
Placebo
DMY
Pancreas
Pancreas
SA-β-Gal positive area (%)
d
WBI
WT
Placebo
DMY
Merge
p16Ink4a
IL6
Kidney
e
p16Ink4a
IL6
Mean fluorescence intensity
f
Kidney
WT
WBI/Placebo
WBI/DMY
Log2 fold change
g
Liver
h
Pancreas
p16Ink4a
p21Cip1
Tnf-α
Il6
Il1β
Tgf-β
Csf2
Cxcl1
Cxcl2
Mmp3
Mmp9

### Supplementary Fig. 12

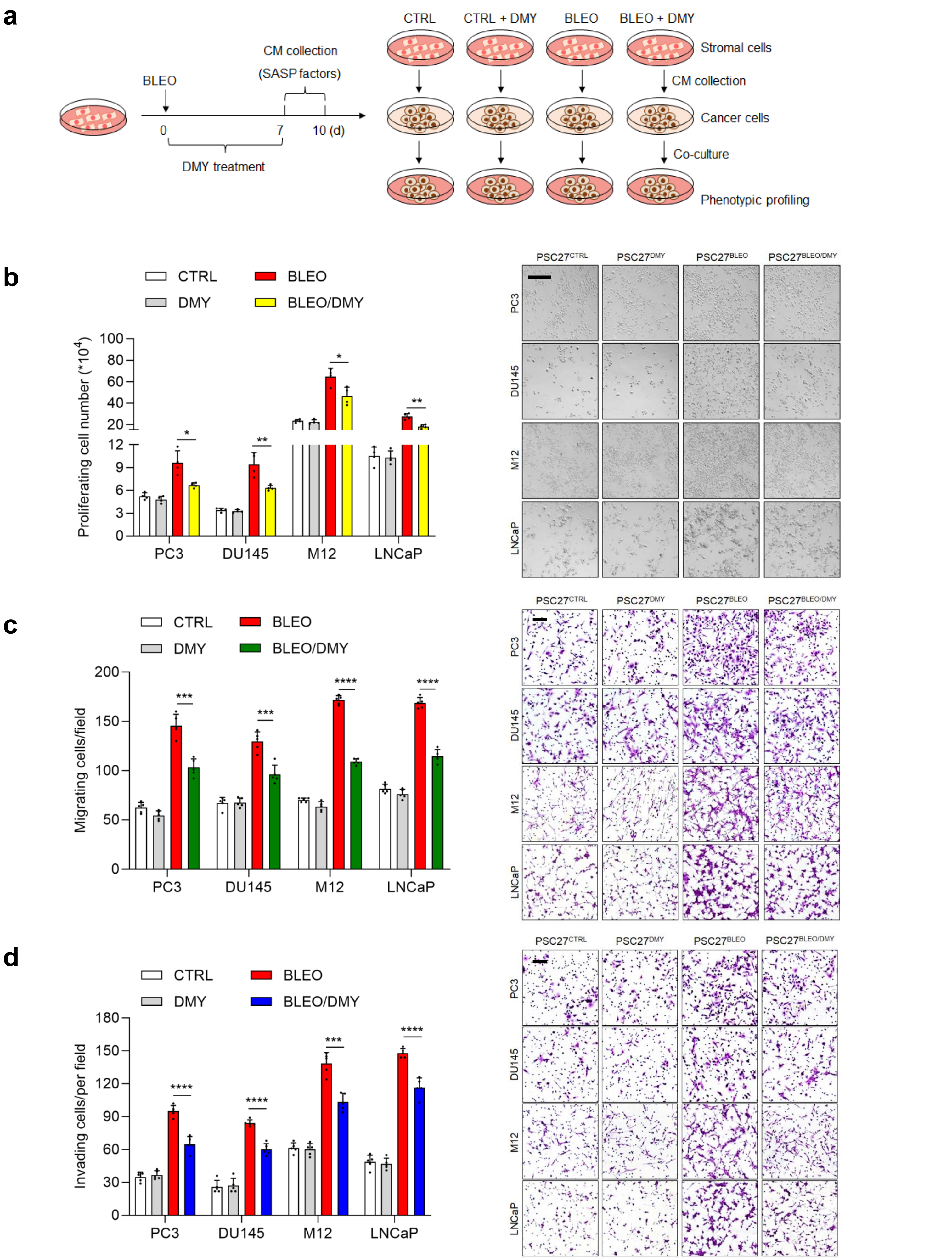

#### Supplementary Fig. 13

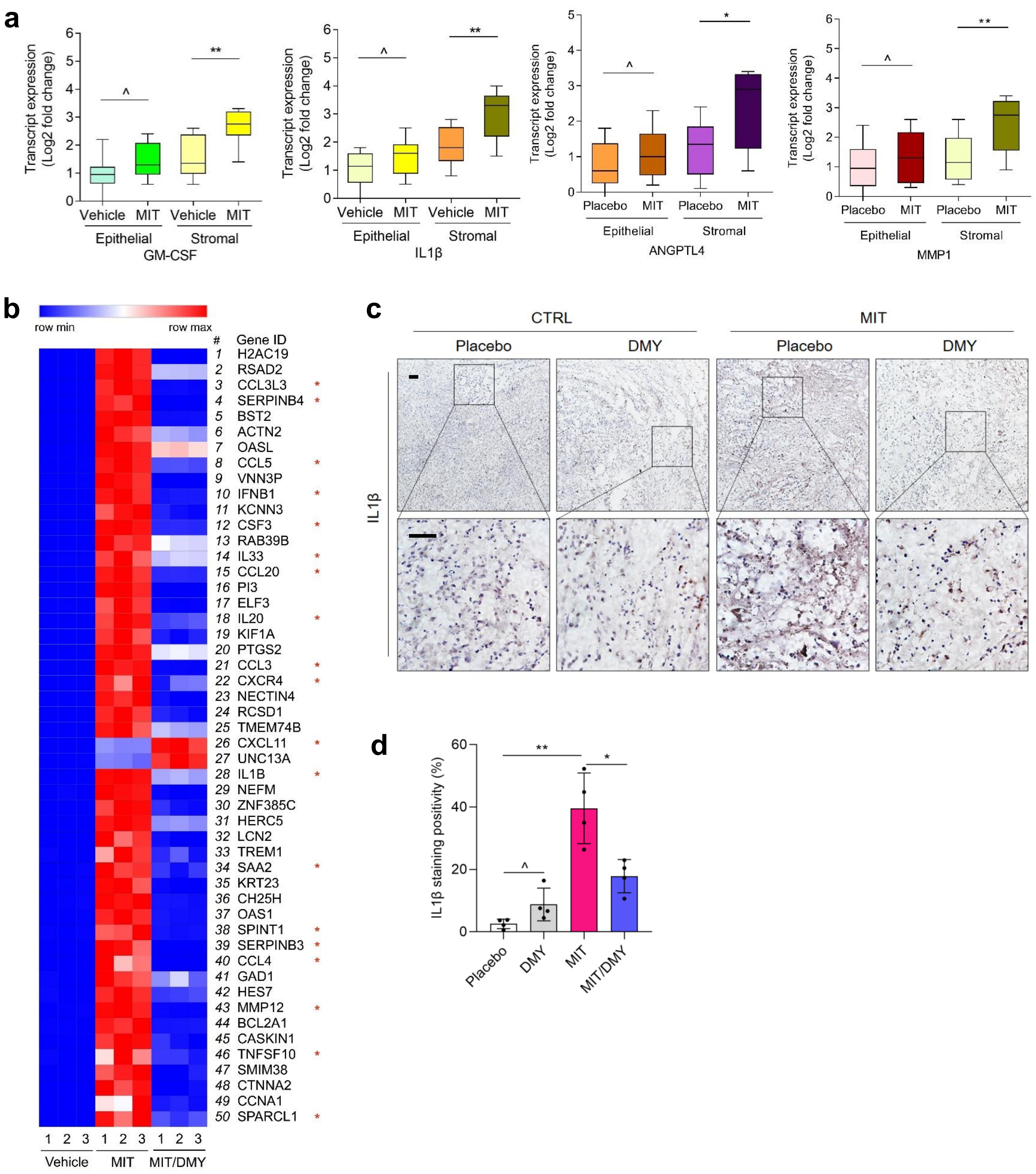

### Supplementary Fig. 14

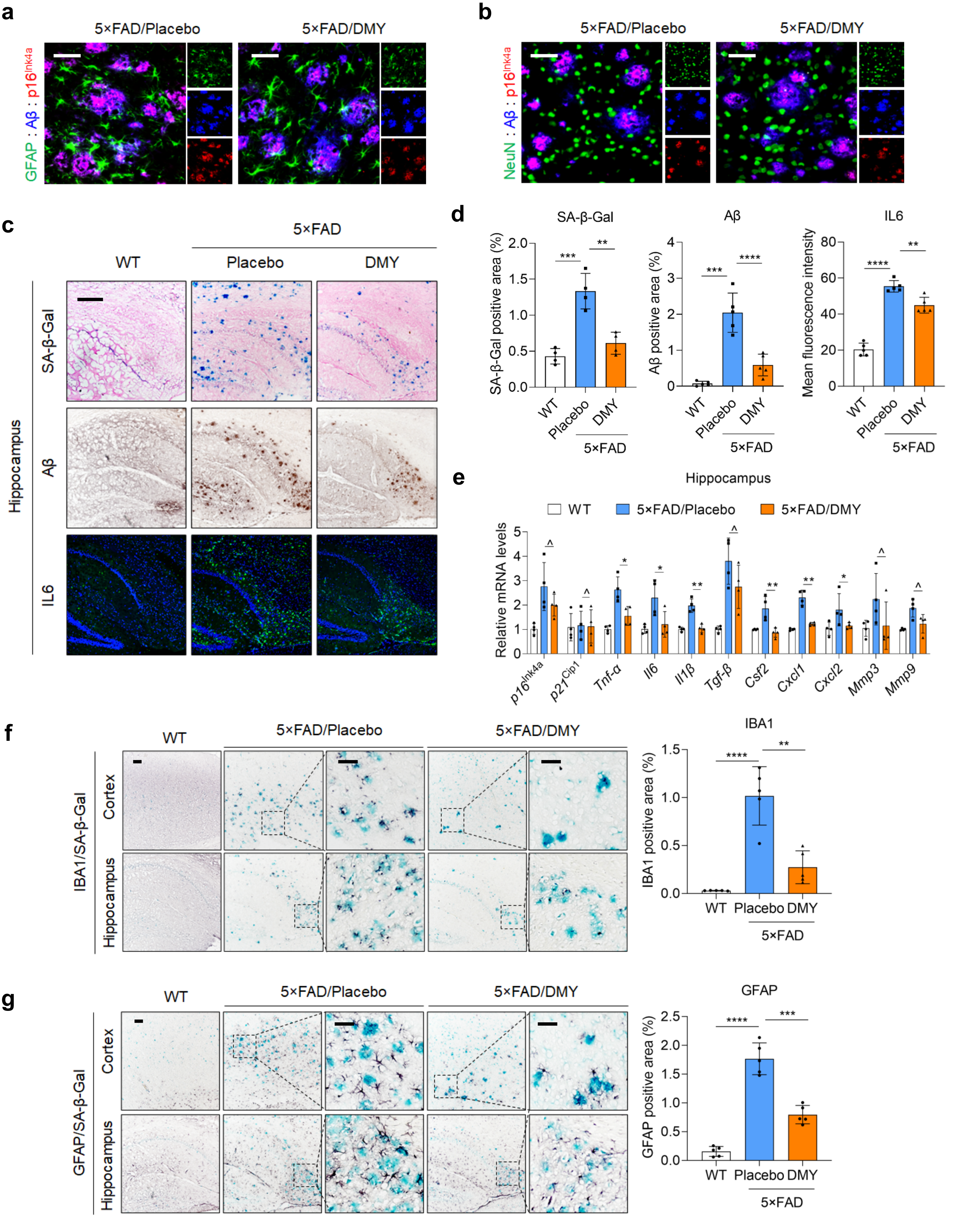
